# Comparative genomics of three non-hematophagous leeches (*Whitmania* spp.): focusing on antithrombotic genes

**DOI:** 10.1101/2024.05.08.590400

**Authors:** Fang Zhao, Zuhao Huang, Lizhou Tang, Bo He, Zichao Liu, Gonghua Lin

**Affiliations:** School of Life Sciences, Jinggangshan University, Ji’an 343009, China; College of Life Sciences, Jiangxi Normal University, Nanchang 330022, China; Engineering Research Center for Exploitation and Utilization of Leech Resources in Universities of Yunnan Province, School of Agronomy and Life Sciences, Kunming University, Kunming 650214, China

**Keywords:** *Whitmania* species, chromosome-level genome, antithrombotic gene, genetic variation, gene expression

## Abstract

Leeches are well known for their blood-feeding habits and are widely used for medicinal purposes as they secrete various antithrombotic substances. However, some leeches such as *Whitmania* spp. exhibit non-hematophagous feeding habits and their significance for medicinal use is debated. In this study, we provide chromosome-level genomes of two non-hematophagous leeches *Whitmania acranulata* and *Whitmania laevis*, and combined with our previous results of *Whitmania pigra*, we systematically analyzed the similarities and differences on the genomes and especially their antithrombotic genes among the three non-hematophagous *Whitmania* leeches. For *W. acranulata*, *W. laevis*, and *W. pigra*, the genome size (181.72 Mb, 173.87 Mb, and 173.56), the percentage of repeat sites (29.55%, 28.28%, and 27.02%), and the number of protein-coding genes (27,068, 23,805, and 24,156) were close to each other, respectively. In contrast, both the total number of the antithrombotic genes (100, 63, and 79), and the detailed constitutes of different antithrombotic gene families were obviously different among the three leeches. There were also massive genetic variations among the members within each antithrombotic gene/protein family. RNA-Seq-based gene expression estimation showed that the expression profiles of the antithrombotic gene families were apparently different among the three leeches. This is the most comprehensive comparison of the genomes and antithrombic biomacromolecules for the *Whitmania* leeches to date. Our results will greatly facilitate the evolutionary research and application of leech derivatives for medical and pharmaceutical purposes of thrombosis.

## Introduction

According to the World Health Organization, cardiovascular diseases directly or indirectly related to thrombophilia claim more than 20 million lives worldwide each year, accounting for 1/3 of all deaths worldwide (WHO 2020; Mensah et al. 2023). Thrombosis is a common complication of cardiovascular disease. Thrombosis occurs when blood clots form in the veins and arteries, blocking the flow of blood and oxygen to the heart and other organs. This can lead to serious health problems, including heart attacks and strokes (DeLoughery 2019). Pharmaceutical institutions have developed a large number of drugs to prevent and treat thrombosis, mainly including anticoagulation, anti-platelet aggregation, and fibrinolysis. Anticoagulants include warfarin, heparin sodium, argatroban, rivaroxaban; antiplatelet agents include aspirin, clopidogrel, and abciximab; and fibrinolytics include streptokinase, alteplase, and ralteplase (Mackman et al. 2020).

Although available antithrombotic drugs have slowed the occurrence of thrombotic events in patients, they have had limited success in reducing the lethality of thrombotic disease. The main reason is that these drugs generally rely on a single target of action and it is difficult to account for individual differences in dosing, which ultimately leads to frequent drug resistance, internal bleeding, liver and kidney damage, and other serious side effects that endanger patients’ lives. For example, the anticoagulant warfarin can cause side effects such as cerebral microbleeds, hemorrhagic stroke and subarachnoid hemorrhage, and leukocyte rupture vasculitis (Elantably et al. 2020; Cheng et al. 2021); the antiplatelet drug clopidogrel carries the risk of drug resistance and increased all-cause mortality (Al-Husein et al. 2018); the fibrinolytic drugs streptokinase and ralteplase, on the other hand, have the potential to induce gastrointestinal bleeding (Giahchi and Mohammadi 2019). Therefore, research and development of safe and effective multi-target drugs with few side effects is an important direction for the treatment of thrombotic diseases.

Leeches (Hirudinea, Annelida) are well known as obligate blood-feeding annelids distributed on all continents except Antarctica (Sawyer 1986). More than 800 species of leeches have been identified (Kvist et al. 2013), some of which are classified as medicinal leeches (Sket and Trontelj 2008). Many leeches feed on mammalian blood, and to ensure smooth satiation within tens of minutes, leeches must secrete a range of biologically active substances such as anticoagulants, thrombolytics, and anti-inflammatories to counteract the host’s physiological hemostatic process (Sig et al. 2017). These bioactive substances, if properly harnessed, can play an important role in medical and medicinal value (Ma et al. 2021). Medicinal leeches, especially the European medical leech (*Hirudo medicinalis*), have been widely used in European countries since the 17th century to treat various diseases such as osteoarthritis (Shakouri et al. 2018), chronic pain (Hohmann et al. 2018), cutaneous leishmaniasis (Hamidizadeh et al. 2017), and postoperative thrombosis prophylaxis (Michalsen et al. 2007; Elyassi et al. 2013). Also, the dried leeches of *Hirudo nipponia* are widely used (orally administered in clinical practice) as traditional Chinese medicine to treat various diseases, especially thrombosis-related diseases such as stroke and coronary heart disease (Dong et al. 2016; Song et al. 2018; Ma et al. 2021).

There are many antithrombotic active components in leeches (Zaidi et al. 2011; Sig et al., 2017), the one that has received the most attention is hirudin. Hirudin has a molecular weight of approximately 7000 daltons and contains approximately 65 amino acid sites (Fritsma 2012; Müller et al. 2020). Hirudin is the most potent natural thrombin inhibitor found to date, binding to thrombin and forming an extremely stable non-covalent complex (Mousa et al. 2021; Montinari and Minelli 2022). Compared to anticoagulants such as warfarin and heparin, hirudin causes fewer bleeding side effects (Chen et al. 2021). In addition to hirudin, leeches have many antithrombotic agents in their bodies and secreted saliva. For example, bdellin, similar to hirudin, showed anticoagulant effects by prolonging the activated partial thromboplastin time (Cheng et al. 2019). Destabilase (Zavalova et al. 1996), leech carboxypeptidase inhibitor (LCI) (Reverter et al. 2000), and hyaluronidase (Jin et al. 2014) were suggested to have thrombolytic properties. Saratin and leech antiplatelet protein (LAPP) were reported to have anti-platelet aggregation function (Gronwald 2008). Also, antistasin and eglin C were thought to have analgesic and anti-inflammatory properties and may indirectly act in thrombophilia (Kvist et al. 2013; Iwama et al. 2020). To our knowledge, two dozen leech-derived antithrombotic compounds have been isolated, involving nearly one hundred gene loci (Babenko et al. 2020; Kvist et al. 2020; Zheng et al. 2021).

Although sanguivory is the most prominent aspect of leech behavior in certain species, other feeding modes such as macrophagy and omnivory are also used by the remaining species (Borda and Siddall 2004). Interestingly, the highly specialized sanguivorous life history mode is not conserved and can be lost repeatedly during the evolutionary history of leeches (Siddall and Burreson 1995). The ancestral Hirudinidae are thought to have been blood-feeders, while at least two genera (*Whitmania* and *Haemopis*) have evolved into invertebrate predators (Borda and Siddall 2004; Phillips and Siddall 2009). Of note are the three *Whitmania* species, *Whitmania pigra* (Whitman, 1884), *Whitmania laevis* (Baird, 1869), and *Whitmania acranulata* (Whitman, 1886), which are sympatric in drainages in China. *W. pigra* is a specific predator of snails; *W. laevis* has a broader diet including snails and insect larvae; while *W. acranulata* feeds on aquatic earthworms and insect larvae (Yang 1996). Morphological studies showed that *W. pigra* and *W. acranulata* retained tooth plates in their jaws, while no tooth plates were detected in *W. laevis* (Qiao et al. 2013), suggesting that *W. laevis* is more completely degenerated than the other two species.

The Pharmacopoeia of the People’s Republic of China (PPRC) is an essential element of China’s drug laws and regulations, and materials not listed in the PPRC should not, in theory, be used for medicinal purposes (Chinese Pharmacopoeia Commission, 2020). The current edition of the PPRC lists three leech species (*H. nipponia*, *W. pigra*, and *W. acranulata*) as legal materials for “Shuizhi” products. For the hematophagous *H. nipponia*, scientists are convinced of its medicinal value, while for *W. pigra* and *W. acranulata*, there has been a long-standing debate as to whether they can serve as the basic source of “Shuizhi” due to their non-blood-sucking habits (He et al. 2021). Some researchers argue that the antithrombotic capabilities of these species were likely lost during their transition from sanguivory to macrophagy, and should therefore be excluded from the PPRC (Song et al. 1997). Conversely, other researchers argue that these two species should be included in the PPRC due to their anticoagulant properties (Ding et al. 2016; Zhang et al. 2020) and antiplatelet aggregation capabilities (Li et al. 1997; Wang et al. 2019), albeit weaker than those of sanguivorous leeches. In addition, recent studies support that at least one type of hirudin from *W. pigra* has anticoagulant activity (Müller et al. 2022). It should be noted that although *W. laevis* was not considered a legal material for “Shuizhi” products, a previous study showed that this species also had some anticoagulant and antiplatelet aggregation activates (Li et al. 1997).

With the rapid development of high-throughput sequencing technology, functional gene research in leeches has now entered the era of genomics. Genomes of several medicinal leeches, such as *H. medicinalis* (Kvist et al. 2020; Babenko et al. 2020), *Hirudinaria manillensis* (Guan et al. 2020; Zheng et al. 2023), *W. pigra* (Tong et al. 2022; Zheng t al. 2023), and *Hirudo nipponia* (Zheng et al. 2023; actually, *Hirudo tianjinensis*) have been published. Recently, we published the most complete genomes to date of *H. manillensis* (Liu et al. 2023) and *W. pigra* (Liu et al. 2024), from which we systematically identified most of the potential antithrombotic genes. Interestingly and unexpectedly, a total of 79 antithrombotic genes were identified from W. pigra, more than the typical blood-feeding *H. manillensis*, which had only 72 antithrombotic genes. Combined with RNA-Seq-based gene expression analyses, our results showed that the number and expression level of antithrombotic genes of a non-hematophagous leech are not always lower than those of a hematophagous leech (Liu et al. 2024). The unique life history of non-hematophagous leech species provides an excellent opportunity to understand the evolution of antithrombotic-related genes in leeches. Here, we provide a chromosome-scale genome of *W. acranulata* and *W. laevis* from which we identified all potential antithrombotic genes of the two species. Furthermore, in combination with RNA-Seq data, we calculated the expression levels of the antithrombotic genes. Combined with our previous results from *W. pigra* (Liu et al. 2024) as well as *H. manillensis* (Liu et al. 2023), we aim to systematically compare the number and expression of antithrombotic genes among the three non-hematophagous *Whitmania* leeches.

## Results

### Basic information of genome assembly

We first used the ONT reads to perform de novo assembly, then used the Survey reads to polish the raw contigs, and finally used the Hi-C reads to consolidate the polished contigs into chromosome-level scaffolds. For *W. acranulata*, a total of 23.23 Gb of ONT reads (N50 length of 24.44 Kb), 20.80 Gb of Survey reads, and 19.33 Gb of Hi-C reads were obtained. After assembly, polishing and consolidation, we obtained 17 scaffolds with a total length of 181.72 Mb and an N50 length of 17.18 Mb. The first 11 longest scaffolds were 23.74 ∼ 11.85 Mb in length, while the remaining 6 scaffolds were less than 0.2 Mb each. Based on the highly discontinuous scaffold length distribution (Fig. 1A) and the well-resolved Hi-C maps (Fig. 1B), we infer that the 11 longest scaffolds are predicted chromosomes. These chromosomes had a total length of 181.23 Mbp, representing 99.73% of the total scaffold length. In addition, we used GetOrganelle to assemble the mitochondrial genome based on the Survey reads. A circular complete mitochondrial genome of 14,505 bp length was obtained (Table 1).

**Figure 1.**
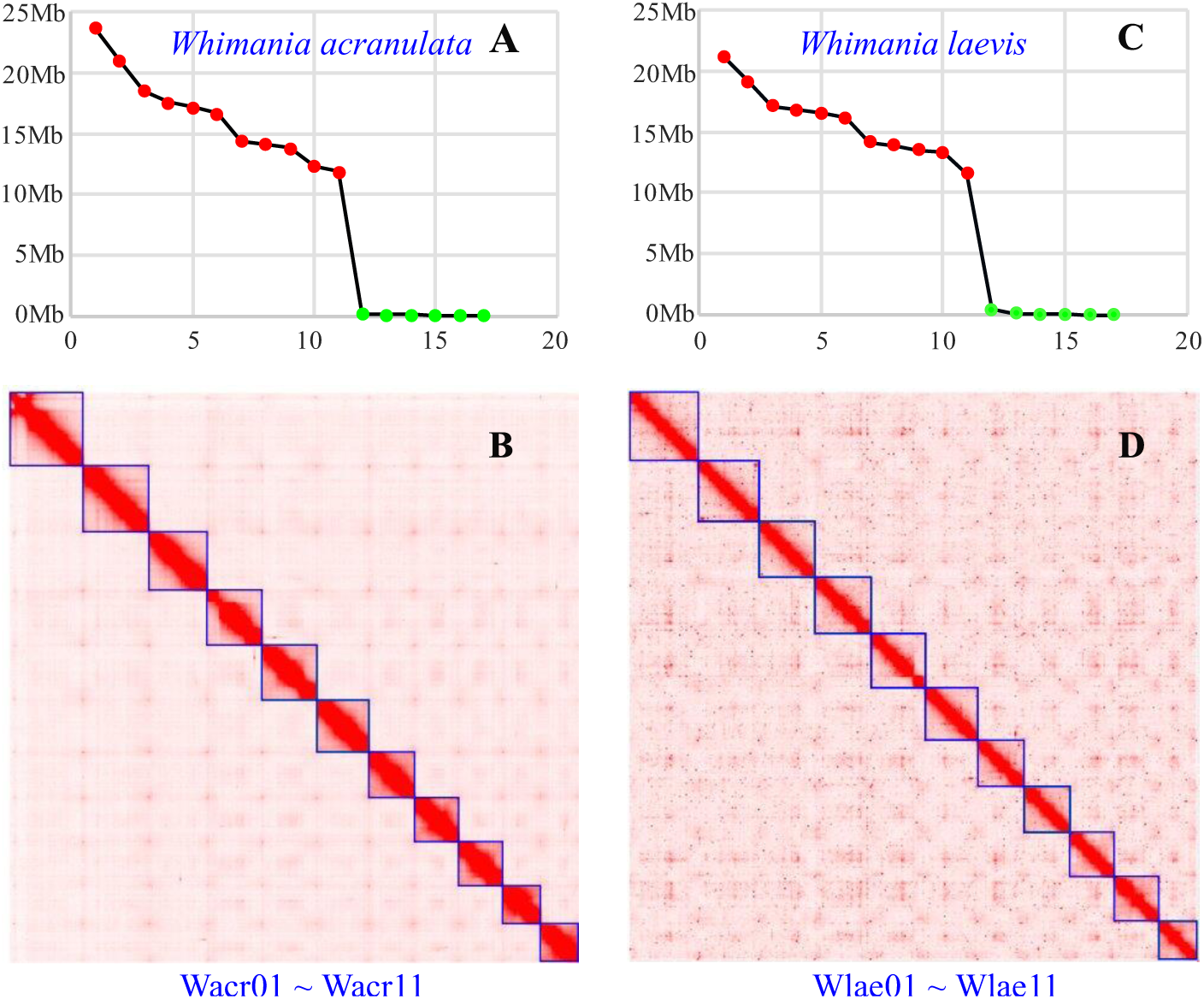
Assembly information of *Whitmania acranulata* and *Whitmania laevis* genomes. (A) Scaffold length distribution of *W. laevis* genome. (B) Chromosome length distribution of *W. laevis* genome. (C) Scaffold length distribution of *W. acranulata* genome. (D) Chromosome length distribution of *W. acranulata* genome.

**Table 1.**
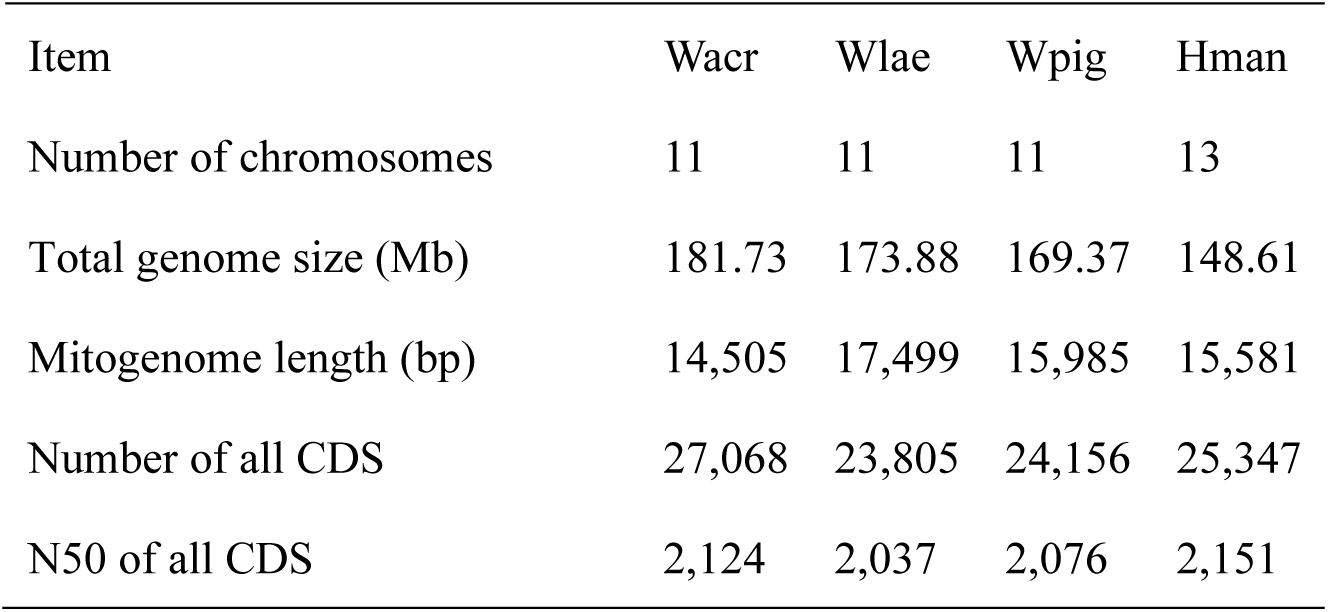
Basic information of genomes and coding sequences (CDS) of *W. acranulata* (Wacr), *W. laevis* (Wlae), *W. pigra* (Wpig) (from Liu et al. 2024), and *H. manillensis* (Hman) (from Liu et al. 2023).

| Item | Wacr | Wlae | Wpig | Hman |
| --- | --- | --- | --- | --- |
| Number of chromosomes | 11 | 11 | 11 | 13 |
| Total genome size (Mb) | 181.73 | 173.88 | 169.37 | 148.61 |
| Mitogenome length (bp) | 14,505 | 17,499 | 15,985 | 15,581 |
| Number of all CDS | 27,068 | 23,805 | 24,156 | 25,347 |
| N50 of all CDS | 2,124 | 2,037 | 2,076 | 2,151 |

As to *W. laevis*, a total of 19.53 Gb of ONT reads (N50 length of 29.78 Kb), 24.38 Gb of Survey reads, and 43.38 Gb of Hi-C reads were obtained. We also obtained 17 scaffolds with a total length of 173.87 Mb and an N50 length of 16.54 Mb. The first 11 longest scaffolds were 21.12 ∼ 11.58 Mb in length, while the remaining 6 scaffolds were less than 0.5 Mb each. The highly discontinuous scaffold length distribution (Fig. 1C) and the well-resolved Hi-C maps (Fig. 1D) also indicated that the species had 11 chromosomes, which had a total length of 173.19 Mbp, representing 99.61% of the total scaffold length. The complete mitochondrial genome assembled from the Survey reads was 17,499 bp in length (Table 1).

The genomes (including all nuclear scaffolds and the mitogenome) of *W. acranulata* and *W. laevis* are available in Supplementary File S1 and File S2, respectively. Combined with our previous study on *H. manillensis* (Liu et al. 2023) and *W. pigra* (Liu et al. 2024), the genome sizes of the three *Whitmania* species were much larger than that of *H. manillensis*, but the other basic information such as mitogenome length and total number of all coding sequences (CDS) were similar among the three *Whitmania* species as well as *H. manillensis* (Table 1).

### Genome quality and repeat sequences

We estimated the completeness of the final assemblies using BUSCO (Benchmarking Universal Single-Copy Orthologs) v5.4.3 (Seppey et al. 2019), with the database eukaryota_odb10. The results showed that, for *W. acranulata*, 250 (98.0%) of the 255 BUSCOs were captured, including 234 (91.8%) complete and single-copy BUSCOs, 8 (3.1%) complete and duplicated BUSCOs, and 8 (1.6%) fragmented BUSCOs; while only 5 (2.0%) BUSCOs were missed. For *W. laevis*, 248 (97.3%) of the 255 BUSCOs were captured, including 236 (92.5%) complete and single-copy BUSCOs, 8 (3.1%) complete and duplicated BUSCOs, and 4 (1.6%) fragmented BUSCOs; while only 7 (2.8%) BUSCOs were missed. We also used Merqury (Rhie et al. 2020) to assess the quality of our genome assembly and obtained quality value scores of 36.54 and 39.92 for the *W. acranulata* and *W. laevis* genomes, respectively.

We searched the genomes for repeat sequences using RepeatModeler v2.0.3 and RepeatMasker v4.1.2-pl (Flynn et al. 2020). A total of 29.55% sites were identified as repeats in the *W. acranulata* genome: retroelements had the highest percentage (12.54%), followed by Unclassified repeats (9.75%) and DNA transposons (5.17%). As to *W. laevis*, a total of 28.28% of the sites were identified as repeats: Unclassified repeats had the highest percentage (10.66%), followed by Unclassified repeats (9.77%) and DNA transposons (6.82%). Combining with our previous study on *H. manillensis* (Liu et al. 2023) and *W. pigra* (Liu et al. 2024), the total percentage of repeat sites of the three *Whitmania* species was much higher than that of *H. manillensis* (Table 2).

**Table 2.** Percentage of different repeat sequence types in the genomes of *W. acranulata* (Wacr), *W. laevis* (Wlae), *W. pigra* (Wpig) (from Liu et al. 2024), and *H. manillensis* (Hman) (from Liu et al. 2023).

| Item | Wacr | Wlae | Wpig | Hman |
| --- | --- | --- | --- | --- |
| Retroelements | 12.54 | 9.77 | 7.82 | 6.59 |
| DNA transposons | 5.17 | 6.82 | 6.80 | 1.76 |
| Rolling-circles | 0.14 | 0.22 | 0.24 | 0.20 |
| Unclassified | 9.75 | 10.66 | 9.46 | 8.41 |
| Small RNA | 0.00 | 0.00 | 0.00 | 0.00 |
| Satellites | 0.00 | 0.15 | 0.00 | 0.20 |
| Simple repeats | 1.95 | 0.66 | 2.70 | 1.80 |
| Low complexity | 0.00 | 0.00 | 0.00 | 0.00 |
| Total | 29.55 | 28.28 | 27.02 | 18.96 |

### Phylogenomics and chromosome syntenic

A total of 766 orthologs were identified among the 11 leech species. Phylogenetic analysis based on the concatenated sequences produced a highly confident consensus tree, i.e. most of the nodes had a bootstrap percentage of 100%. As expected, the three *Whitmania* species and the three *Hirudo* species respectively formed two monophyletic clades, which were the most closely related among the genera in the tree. *W. pigra* and *W. laevis* were more closely related to each other than *W. acranulata* (Figure 2A).

**Figure 2.**
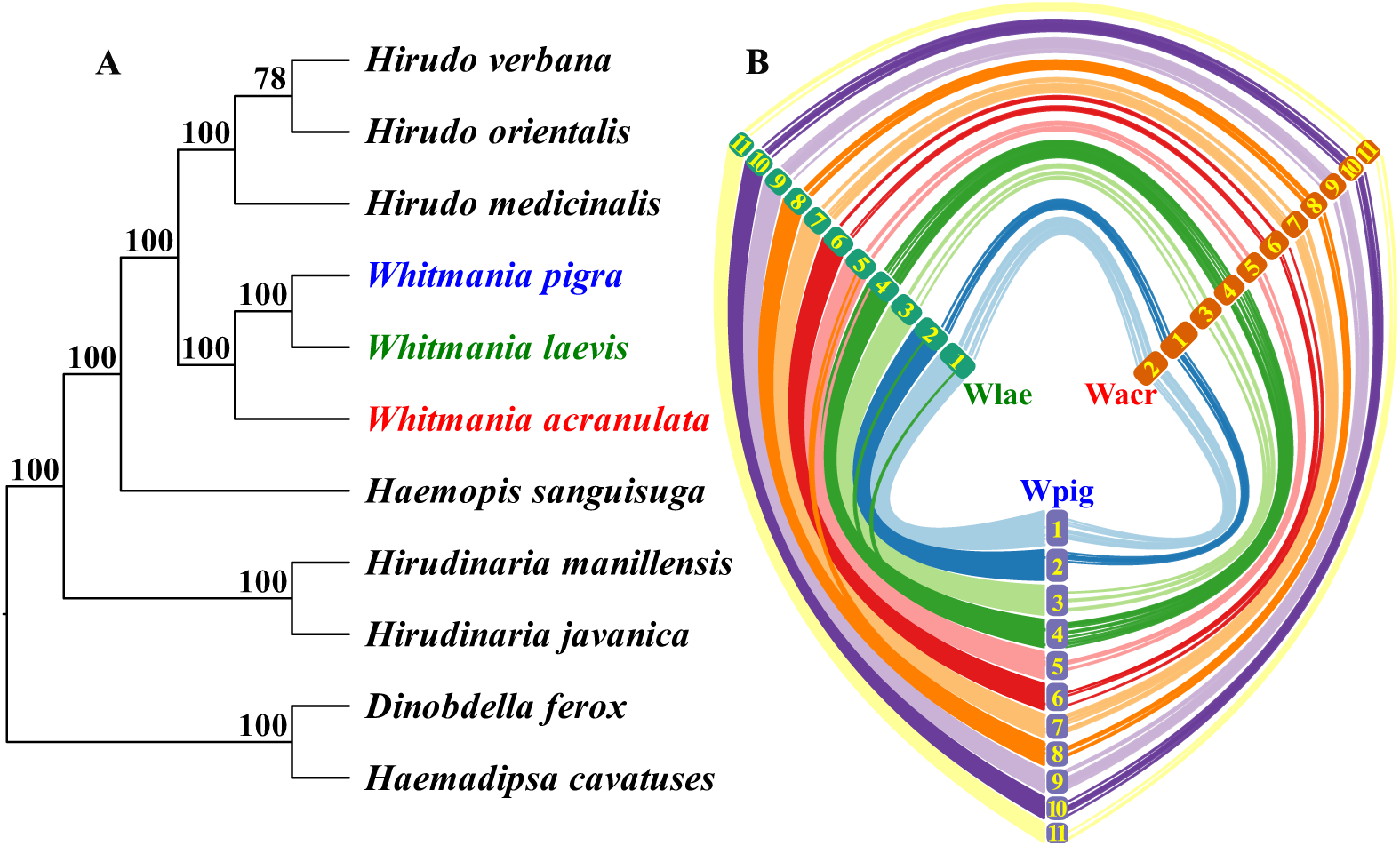
Genomic relationships among the three *Whitmania* leeches. (**A**) Phylogenomics of the three Whitmania species as well as other related taxa; (**B**) Chromosome syntenic relationships among the three species (more curves indicate higher collinearity levels; the numbers indicate the chromosomes of each species; larger numbers represent smaller chromosomes).

Chromosome syntenic analysis showed that the chromosomes of *W. pigra* and *W. laevis* were perfectly matched. The chromosome length order is identical and there were massive syntenic segments between the two species. In contrast, there was an inconsistency in chromosome length order between *W. acranulata* and the other two *Whitmania* species, i.e. the longest chromosome (Wacr01) of *W. acranulata* corresponded to the second longest chromosome of *W. pigra* (Wpig02) and *W. laevis* (Wlae02), while the second longest chromosome (Wacr02) corresponded to the longest chromosome of *W. pigra* (Wpig01) and *W. laevis* (Wlae01). Furthermore, the syntenic segments between the *W. acranulata* and the other two *Whitmania* species were obviously fewer than those between *W. pigra* and *W. laevis* (Figure 2B).

### Constitution of antithrombotic genes

Based on the BRAKER-plus strategy, 27,068 protein-coding genes with a total length of 39,864,801 bp and an N50 length of 2,124 bp were predicted for *W. acranulata*. For *W. laevis*, 23,805 protein-coding genes were predicted, with a total length of 34,232,948 bp and an N50 length of 2,037 bp. The GFF file and all predicted CDSs of the two species (Supplementary File S3 ∼ S6) were available as Supplementary Material files. A total of 100 and 63 antithrombotic genes were identified from the genomes of *W. acranulata* and *W. laevis*, respectively. According to our previous study on *W. pigra* (Liu et al. 2023; Liu et al. 2024), these genes could be classified into different gene families (Table 3; for more information on the functions of these protein families, see also the two previous papers and the references therein).

**Table 3.** Gene numbers of the antithrombotic gene families of *W. acranulata* (Wcar), *W. laevis* (Wlae), *W. pigra* (Wpig) and *H. manillensis* (Hman).

| Gene family | Wacr | Wlae | Wpig | Hman | Function |
| --- | --- | --- | --- | --- | --- |
| *hirudin* | 0 | 0 | 7 | 5 | coagulation inhibitor |
| *progranulin* | 1 | 1 | 1 | 1 | coagulation inhibitor |
| *antistasin* | 3 | 3 | 2 | 2 | coagulation inhibitor |
| *lefaxin* | 3 | 3 | 3 | 3 | coagulation inhibitor |
| *therostasin* | 1 | 1<sup>#</sup> | 1 | 1 | coagulation inhibitor |
| *hirustasin* | 1 | 2 | 1 | 1 | coagulation inhibitor |
| *hirustasin-like* | 8 | 8 | 9 | 12 | coagulation inhibitor |
| *guamerin* | 1 | 1 | 1 | 1 | coagulation inhibitor |
| *piguamerin* | 3 | 1 | 1<sup>#</sup> | 1 | coagulation inhibitor |
| *bdellastasin* | 1 | 1 | 1 | 1 | coagulation inhibitor |
| *poecistasin* | 0 | 0 | 0 | 2 | coagulation inhibitor |
| *eglin* | 11 | 5 | 2 | 4 | coagulation inhibitor |
| *bdellin* | 3 | 2 | 1 | 1 | coagulation inhibitor |
| *LDTI* | 4 | 1 | 1 | 1 | coagulation inhibitor |
| *HMEI* | 35<sup>##</sup> | 8 | 20<sup>#</sup> | 18 | coagulation inhibitor |
| *saratin* | 0 | 6<sup>#</sup> | 11 | 2 | platelet aggregation inhibitor |
| *apyrase* | 9 | 4 | 3 | 5 | platelet aggregation inhibitor |
| *lumbrokinase* | 2 | 4 | 4 | 3<sup>#</sup> | platelet aggregation inhibitor |
| *destabilase* | 6 | 6 | 4 | 3<sup>#</sup> | fibrinolysis enhancer |
| *GGT* | 4 | 2 | 2<sup>#</sup> | 1 | fibrinolysis enhancer |
| *LCI* | 1 | 1 | 1 | 1 | fibrinolysis enhancer |
| *hyaluronidase* | 3 | 3 | 3 | 3 | tissue penetration enhancer |
| total | 100 | 63 | 79 | 72 | — |

LDTI, leech-derived tryptase inhibitor; *Hirudinaria manillensis* elastase inhibitor; GGT, gamma-glutamyl transpeptidase; LCI, leech carboxypeptidase inhibitor; <sup>#</sup>, including one pseudogene; <sup>##</sup>, including two pseudogenes; —, Not applicable.

Unlike *W. pigra*, which had two thrombin inhibitor families (*hirudin* and *progranulin*), only *progranulin* was identified in *W. acranulata* and *W. laevis*. Three factor Xa inhibitor families (*antistasin*, *lefaxin*, and *therostasin*) were recovered in both *W. acranulata* and *W. laevis*. As in *W. pigra*, most of the remaining coagulation inhibitors (*hirustasin*, *hirustasin-like*, *guamerin*, *piguamerin*, *bdellastasin*, *eglin*, *bdellin*, *LDTI*, and *HMEI*) were recovered in both *W. acranulata* and *W. laevis*. Of the three antiplatelet families (*saratin*, *apyrase*, and *lumbrokinase*), two and three families were recovered in *W. acranulata* and *W. laevis*, respectively. Finally, three families of fibrinolysis enhancers (*lumbrokinase*, *destabilase*, *GGT*, and *LCI*) and one family of tissue penetration enhancers (hyaluronidase) were also recovered in both *W. acranulata* and *W. laevis* (Table 3).

### Variation of antithrombotic genes/proteins

There was massive genetic variation among the members in each of the antithrombotic gene/protein families (Figure S1 ∼ S17), including many pseudogenes (Table2). Although hirudin is the first identified and most representative antithrombotic bioactive molecule in leeches (Krezel et al. 1994), the genes were completely lost from the genomes of *W. acranulata* and *W. laevis* (Figure S1), making it impossible to show their interspecific genetic variation. One or more members of the gene families *therostasin* (*therostasin_Wlae*, Figure S5), *piguamerin* (*piguamerin_Wpig*, Figure S6), *HMEI* (*HMEI_Wacr25*, *HMEI_Wacr28*, *HMEI_Wpig07*; Figure S10), *saratin* (*saratin_Wlae4* and *saratin_Wpig03*, Figure S11), and *GGT* (*GGT_Wpig2*, Figure S15).

There were many proteins whose reactive residues were predicted by functional verification tests. The arginine from the C**R**EHC segment was predicted to be the catalytic residue in the archetypal antistasin (Dunwiddie et al. 1989), but of the eight proteins from the three *Whitmania* species, only one (antistasin_Wpig2) was conserved at this site (Figure S3). The predicted catalytic residue arginine (C**R**IYC) in the archetypal therostasin (Chopin et al. 2000) was replaced by phenylalanine in all therostasins from the three *Whitmania* species (Figure S5). The catalytic arginine (C**R**IRC) of the archetypal hirustasin (Mittl et al. 1997) was conserved in the hirustasins of *W. pigra*, but not in those of the other species (Figure S6). The catalytic arginine (C**R**KYC) of the archetypal piguamerin (Kim and Kang 1998) was conserved in all piguamerins except piguamerin_Wlae (Figure S6). The catalytic methionine (C**M**IFC) of the archetypal guamerin (Jung et al. 1995) and the catalytic arginine (C**K**VKC) of the archetypal bdellastasin (Moser et al. 1998) were conserved in all guamerins and bdellastasins from the three *Whitmania* species (Figure S6). Futhermore, the corresponding coding sequences of the two residues were also conserved in the three *Whitmania* species (Figure 4), suggesting that the two gene families may also be functionally conserved in the *Whitmania* species.

The reactive residues leucine and asparagine (McPhalen et al. 1985) in the archetypal eglin (T**LD**LR) were conserved in 7 of 15 eglins from the three *Whitmania* species (Figure S7). The reactive residue lysine (CT**K**EL) in the archetypal bdellin (Kim et al. 2001) was conserved in three of five bdellins from the *Whitmania* species (Figure S8). The reactive residues lysine and isoleucine (P**KI**LK) of the archetypal LDTI (Sommerhoff et al. 1994) were conserved in the LDTIs of *W. laevis* and *W. pigra*, but not those of *W. acranulata* (Figure S9). The catalytic histidine (KI**H**NM) of the archetypal destabilase (Marin et al. 2023) was conserved in all destabilases from the three *Whitmania* species, except for destabilase_Wlae5 (Figure S14). Finally, the catalytic threonine (HG**T**AH) of the archetypal GGT (Castellano et al. 2012) was conserved in all GGTs from the three *Whitmania* species (Figure S15).

### Expression of antithrombotic genes

Based on the RNA-Seq data sequenced in this study we calculated the total TPM of each antithrombotic gene family of the three *Whitmania* species. We then performed pairwise comparisons on the TPM of each gene family among the three species using the non-parametric Mann-Whitney U test. The results showed that, of the 21 gene families, 17 were significantly different between at least two species (Table 4). Three gene families (*hirudin*, *antistasin*, and *therostasin*) in *W. pigra* had higher expression levels than the other two *Whitmania* species; two gene families (*eglin* and *apyrase*) in *W. acranulata* had higher expression levels than the other two *Whitmania* species; five gene families (*hirustasin*, *hirustasin-like*, *guamerin*, *bdellastasin*, and *bdellin*) in *W. laevis* had higher expression levels than the other two *Whitmania* species. Hierarchical cluster analysis showed that the samples were grouped into three clusters, each corresponding to one of the three *Whitmania* species. The *W. pigra* and *W. laevis* samples had closer expression patterns, while the *W. acranulata* samples were less similar to the other two *Whitmania* species (Figure 3).

**Figure 3.**
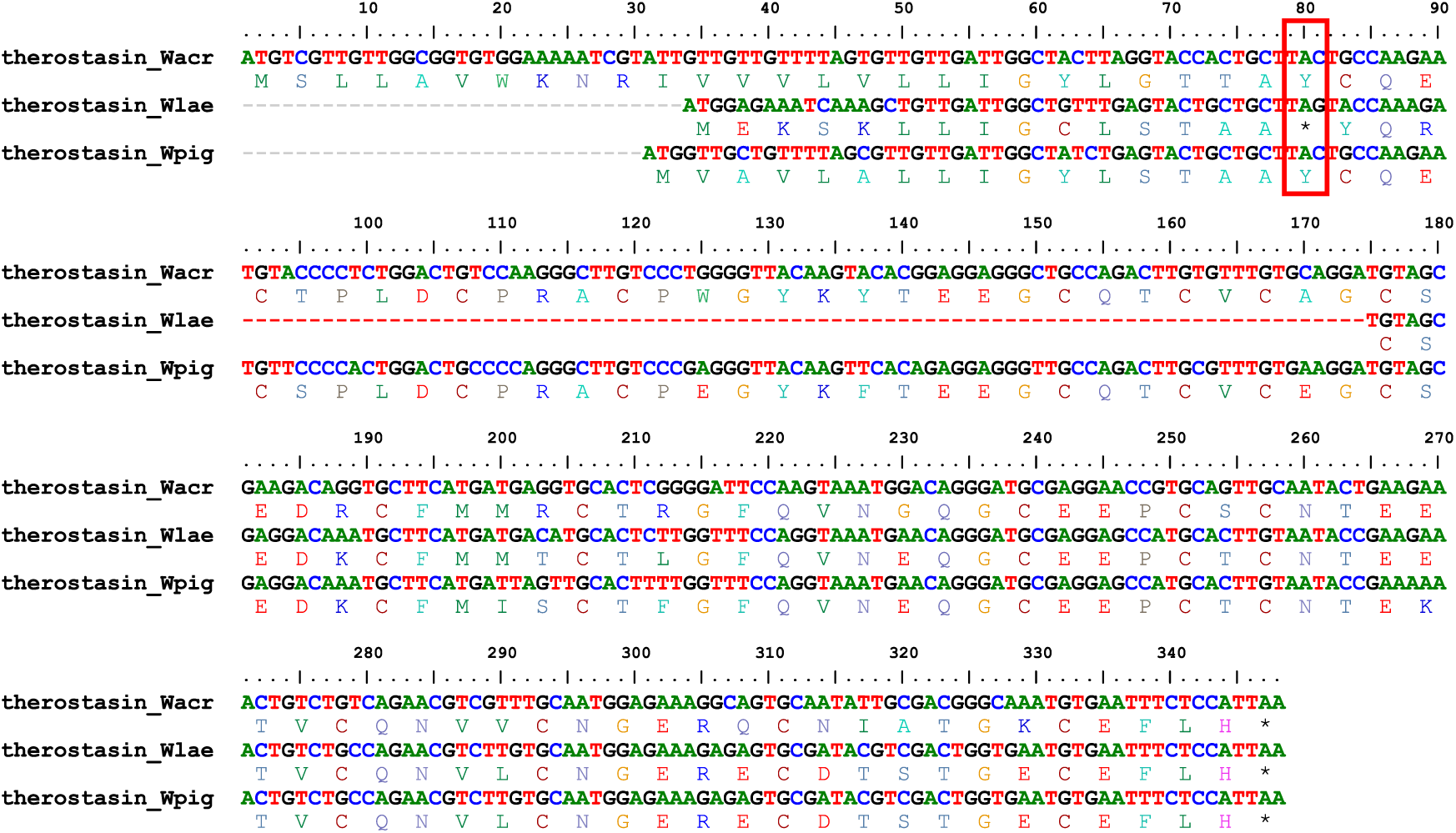
Alignment of therostasins.

**Figure 4.**
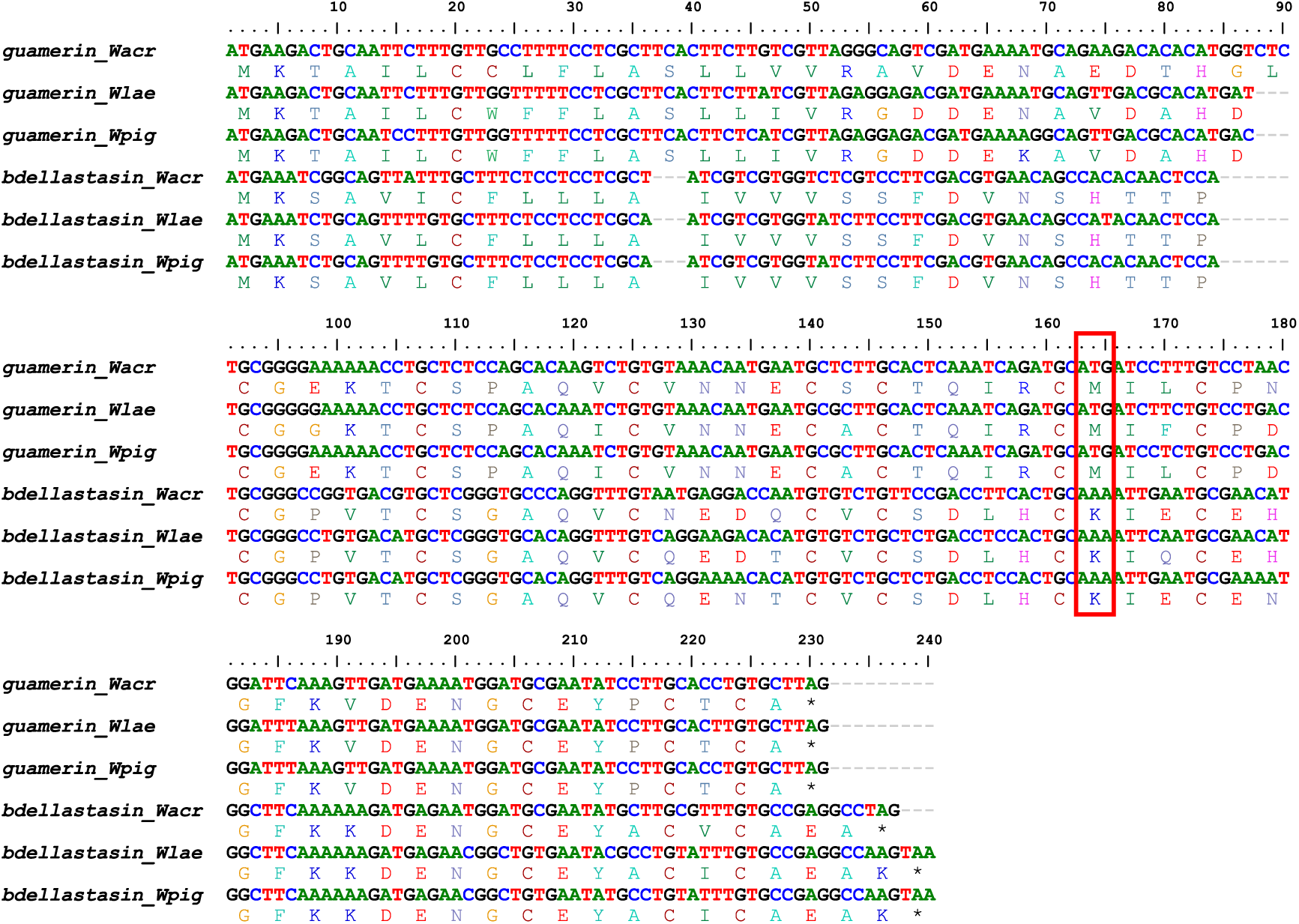
Alignment of antistasins.

**Figure 5.**
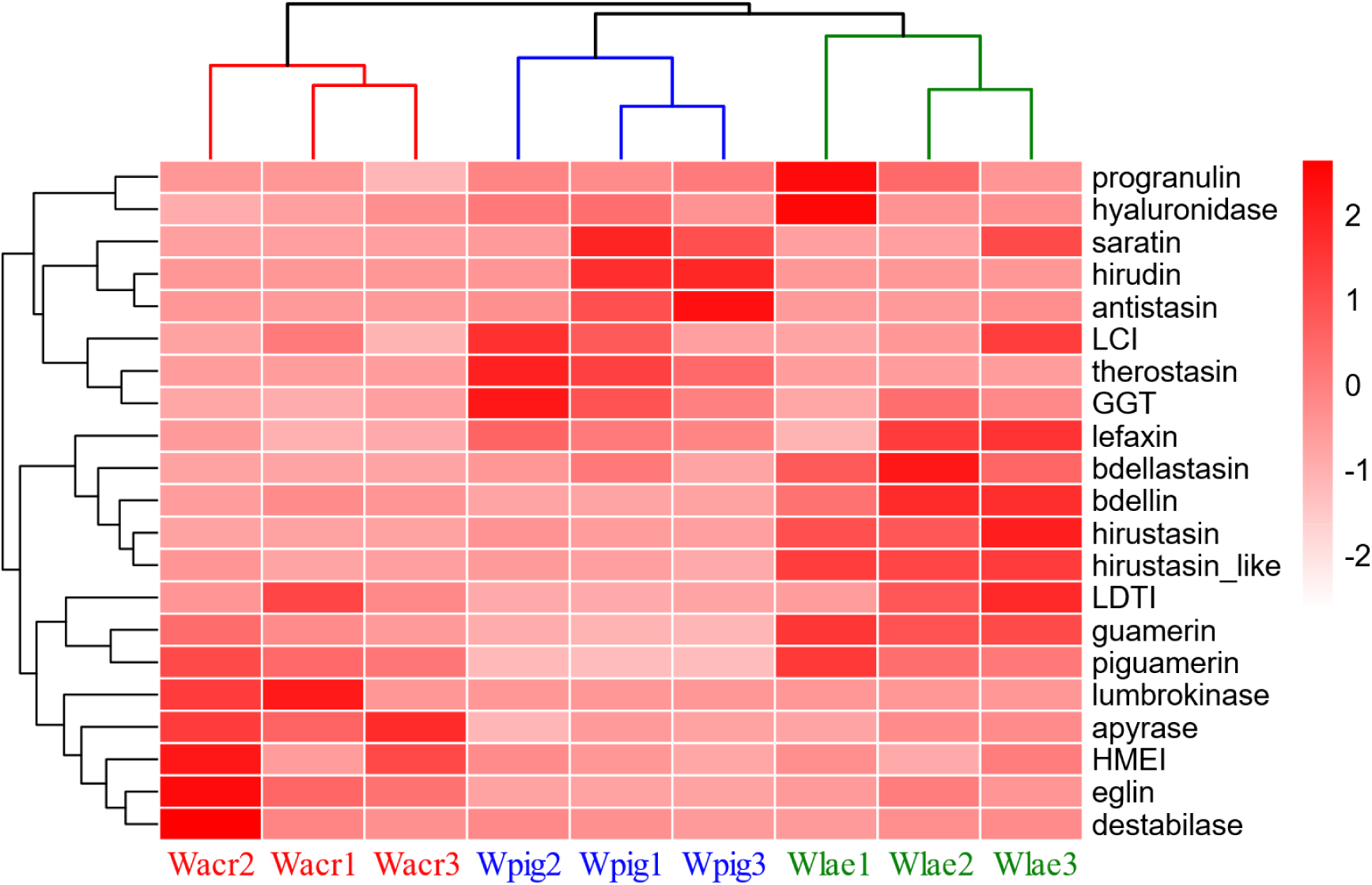
Heatmap of the expression level of antithrombotic genes of the three Whitmania species (*W. pigra*, Wpig1 ∼ 3, SRA No. SRR26541745 ∼ SRR26541743; *W. acranulata*, Wacr1 ∼ 3, SRA No. SRR27841062 ∼ SRR27841060; *W. laevis*, SRA No. Wlae1 ∼ 3, SRR26541749 ∼ SRR26541747).

**Table 4.** Comparison of the total TPM of each gene family among the *Whitmania* samples (Different letters in the upper right indicate significant differences at the level of *P* < 0.05; —, not applicable due to gene family loss).

| Gene family | Wacr | Wlae | Wpig |
| --- | --- | --- | --- |
| *hirudin* | — | — | 11929.3 ± 9848.5<sup>a</sup> |
| *progranulin* | 24.9 ± 20.3<sup>c</sup> | 109.7 ± 82.9<sup>ab</sup> | 61.3 ± 9.5<sup>b</sup> |
| *antistasin* | 163.8 ± 72.5<sup>b</sup> | 617.7 ± 991.2<sup>b</sup> | 11459.8 ± 9897.5<sup>a</sup> |
| *lefaxin* | 4359.6 ± 1049.1<sup>b</sup> | 11109.3 ± 6842.0<sup>ab</sup> | 8970.7 ± 1647.5<sup>a</sup> |
| *therostasin* | 0.1 ± 0.0<sup>b</sup> | 0.3 ± 0.5<sup>b</sup> | 79.6 ± 31.5<sup>a</sup> |
| *hirustasin* | 125.6 ± 29.8<sup>c</sup> | 8306.8 ± 2673.8<sup>a</sup> | 702.3 ± 570.1<sup>b</sup> |
| *hirustasin-like* | 2978.1 ± 1346.0<sup>b</sup> | 19232.3 ± 831.5<sup>a</sup> | 2157.3 ± 1260.2<sup>b</sup> |
| *guamerin* | 8833.9 ± 3894.1<sup>b</sup> | 19477.3 ± 2340.6<sup>a</sup> | 1485.1 ± 855.7<sup>c</sup> |
| *piguamerin* | 18474.4 ± 4611.1<sup>a</sup> | 19208.9 ± 6981.9<sup>a</sup> | 251.2 ± 149.0<sup>b</sup> |
| *bdellastasin* | 255.1 ± 45.1<sup>b</sup> | 2823.1 ± 1193.7<sup>a</sup> | 725.3 ± 626.9<sup>b</sup> |
| *eglin* | 9791.0 ± 6286.7<sup>a</sup> | 2438.7 ± 1602.2<sup>b</sup> | 323.9 ± 100.3<sup>b</sup> |
| *bdellin* | 3152.0 ± 1592.9<sup>b</sup> | 17181.9 ± 6960.1<sup>a</sup> | 620.0 ± 165.5<sup>c</sup> |
| *LDTI* | 884.4 ± 705.3<sup>a</sup> | 1254.7 ± 955.0<sup>a</sup> | 63.4 ± 48.6<sup>b</sup> |
| *HMEI* | 12434.4 ± 6605.3<sup>a</sup> | 6399.2 ± 2160.0<sup>a</sup> | 5608.7 ± 1492.1<sup>a</sup> |
| *saratin* | — | 1888.5 ± 3263.7<sup>a</sup> | 4580.8 ± 4007.4<sup>a</sup> |
| *apyrase* | 61.7 ± 14.5<sup>a</sup> | 22.4 ± 7.4<sup>b</sup> | 12.9 ± 6.7<sup>b</sup> |
| *lumbrokinase* | 4061.2 ± 3625.9<sup>ac</sup> | 38.0 ± 5.5<sup>c</sup> | 15.2 ± 9.9<sup>ab</sup> |
| *destabilase* | 32661.6 ± 36165.4<sup>a</sup> | 8721.2 ± 3313.1<sup>a</sup> | 8672.2 ± 4262.8<sup>a</sup> |
| *GGT* | 46.8 ± 9.1<sup>bc</sup> | 83.7 ± 35.5<sup>ab</sup> | 159.5 ± 67.4<sup>a</sup> |
| *LCI* | 1724.3 ± 585.2<sup>a</sup> | 2329.4 ± 1162.0<sup>a</sup> | 2889.3 ± 1157.1<sup>a</sup> |
| *hyaluronidase* | 29.9 ± 10.6<sup>a</sup> | 69.0 ± 51.7<sup>a</sup> | 50.7 ± 12.2<sup>a</sup> |

## Discussion

### Similarities in the three *Whitmania* genomes

A high-quality genome is essential for molecular evolutionary analyses of organisms. Recent high-throughput sequencing and assembly technologies make it easier to obtain a chromosome-scale genome. Here, we used the third- and the next-generation sequencing methods to obtain chromosome-scale genomes of *W. acranulata* and *W. laevis*. For both species the predicted chromosomes constitute over 99.5% of the total scaffold length, indicating that the genome assemblies have high sequence continuity. The BUSCO analyses showed that over 97% of the BUSCOs were captured in the two genomes, indicating their high completeness. The Merqury analyses yielded quality scores above 36, higher than the previous study on other leech species (Zheng et al. 2023). In addition, we assembled a complete circular mitochondrial genome for each of the two leeches. As a result, based on the above parameters, we have for the first time provided high quality and nearly complete genomes of the two non-hematophagous leeches.

Combing the genomes of *H. manillensis* and *W. pigra* obtained in our previous studies (Liu et al. 2023; Liu et al. 2024), we were able to identify the similarities in the three non-hematophagous *Whitmania* genomes, compared to the hematophagous species *H. manillensis*. First, the three *Whitmania* species had a similar genome size, which was however, obviously larger than that of *H. manillensis*. One of the most important factors contributing to genome size is the presence of repetitive sequences. Different types of repetitive DNA elements such as transposable elements, simple sequence repeats, and others, play a significant role in shaping genome size in different organisms (Liao et al. 2023). The total percentage of repeat sites of the three *Whitmania* species was over 27%, much higher than that of *H. manillensis* (18.96%). The most striking case was that, the percentage of DNA transposons of *Whitmania* species was about three times of that of *H. manillensis*. It is well known that repeat sequences can be a source of genetic variation, influence gene regulation, affect genome stability and structure, and ultimately contribute to the adaptive potential of organisms (Papworth and Pevzner 2018). *Whitmania* species were previously placed in Haemopidae, which was a sister family to Hirudinidae. However, recent studies have shown that *Whitmania* is an ingroup within the genus *Hirudo*, suggesting that the ancestor of *Whitmania* species was a blood-feeder (Phillips and Siddall 2009; Zhao et al. 2021). We hypothesize that the abundance of genetic material may have played an important role in the evolutionary events of diet switching in these *Whitmania* species.

Despite the discrepancy in genome size, the number of protein-coding genes of the *Whitmania* species (25,010 ± 1,791) was close to that of *H. manillensis* (25,347). It should be noted that, of the 22 antithrombotic gene families identified in *H. manillensis*, three (*hirudin*, *poecistasin*, and *saratin*) were lost in one or more *Whitmania* species. It is apparent that the habit changes from hematophagous to non-hematophagous has resulted in the loss of antithrombotic genes due to reduced selection pressure on these genes. In addition, similar to *H. manillensis*, all the three *Whitmania* species had two or more pseudogenes, suggesting that pseudogenization of antithrombotic genes was common in leeches, regardless of their habit.

### Differences among *Whitmania* species

Although, as mentioned above, the *Whitmania* species had many similarities in their genomes and antithrombotic genes, the genetic differences among the three species deserve to be mentioned. First, the constitutions of repeat sequences were different among the three species. For example, the percentage of retroelements in *W. acranulata* was much higher than that in *W. pigra* and *W. laevis*. Also, *W. laevis* had much fewer simple repeats than the other two *Whitmania* species as well as *H. manillensis*. Second, although the phylogenetic analysis confirmed the monophyletic relationships of the three *Whitmania* species, the chromosome syntenic analysis showed that the genome collinearity between *W. acranulata* and the other two *Whitmania* species was significantly weakened. These results suggest that the *Whitmania* species had active genomic variation, which may be related to its richness in repetitive sequences.

The number of antithrombotic genes also differed among the three *Whitmania* species. The most striking feature was that a total of 100 antithrombotic genes were identified in *W. acranulata*, while only 63 were found in *W. laevis*. In the gene family *HMEI*, 35 members were found in *W. acranulata*, while only eight were found in *W. laevis*. In the eglin family, 11 were found in *W. acranulata*, while only two were found in *W. pigra*. Gene loss was also unpredictable, e.g. seven *hirudins* were found in *W. pigra*, but this gene family disappeared completely in *W. acranulata* and *W. laevis*. Similarly, 11 and six saratins were identified in *W. pigra* and *W. laevis*, respectively, whereas this gene family was completely undetectable in the *W. acranulata* genome. Furthermore, there was massive genetic variation among the members of each gene/protein family. Taking the reactive residues as an example, among the 11 protein families, only three (guamerin, bdellastasins, and GGT) had all members sharing the same residues with their corresponding archetypal proteins. These results indicate that both the number and the sequences of antithrombotic genes/proteins are extremely active in *Whitmania* leeches, probably also due to relaxed selection pressure on these families.

The hierarchical cluster analysis based on the TPM of the antithrombotic families showed that the samples from each species perfectly formed their respective clusters, indicating that the expression profiles of these gene families were apparently different among these *Whitmania* species. Consistent with the results of the phylogeny and chromosome synteny analyses, the antithrombotic gene expression profile also showed that *W. laevis* was closer to *W. pigra*, while *W. acranulata* was more distantly related to them. Looking specifically at each gene family, there were many differences in TPM levels among *Whitmania* species. For example, the expression level of five gene families (*hirustasin*, *hirustasin-like*, *guamerin*, *bdellastasin*, and *bdellin*) was significantly higher than that of *W. pigra* and *W. acranulata*. In contrast, the expression level of the *eglin* and *apyrase* families was much higher in *W. acranulata* than in *W. pigra* and *W. laevis*. These results suggest that, different species have evolved different expression regulation of these genes for different survival strategies.

### Perspectives on pharmacological application

Leeches have been used as a medical and pharmaceutical resource for many centuries (Michalsen et al. 2007). In addition to hematophagous species, some non-hematophagous leeches have also been used as traditional Chinese medicine “Shuizhi” for the treatment of antithrombotic diseases (Chinese Pharmacopoeia Commission, 2020; Yu et al. 2022). *W. pigra*, which has the largest body size, the most abundant resources, and is relatively easy to cultivate, has become the primary material for “Shuizhi” (Liu and Yang 2014). In contrast, *W. acranulata* and *W. laevis* were less concerned due to the scarcity of resources. It has been repeatedly confirmed that, all three *Whitmania* species mentioned in this study have anticoagulant and/or antiplatelet activities (Ou et al. 1996; Li et al. 1997; Guan et al. 2012; Zhang et al. 2012; Li et al. 2014; Ding et al. 2016; Cheng et al. 2018; Wang et al. 2019; Zhang et al. 2020; Zhong et al. 2020). It should be noted that, as mentioned above, the three species had considerable deviations in genome characteristics and the constitution and expression patterns on antithrombotic genes, detailed pharmacological characteristics among these species need to be determined in the future.

Probably due to the complexity of the functional activity and expression level of the different antithrombotic genes, there were inconsistencies in different studies on the antithrombotic activities of the *Whitmania* species. For example, Wang et al. (2013) showed that *W. pigra* and *W. acranulata* had similar antithrombin activities, while Cheng (2018) indicated that the antithrombin activities of *W. pigra* were higher than those of *W. acranulata*. In contrast, Zhang et al. (2012) and Zhang et al. (2020) showed that the antithrombin activities of *W. pigra* were lower than that of *W. acranulata*. Even for the same species such as *W. pigra*, different sample sources (wild vs. artificial rearing) (Li et al. 1997) and extraction methods (water extraction vs. bionic extraction) (Tan et al. 2019) may result in significant differences in their antithrombin activities. The large number of antithrombotic genes identified in this study, and the availability of complete CDSs for most of these genes (except for pseudogenes), provided a large opportunity to test their function by producing recombinant proteins (Gupta et al. 2016) and performing in vitro (Liu et al. 2023) and in vivo (Tang et al. 2018) experiments.

Hirudin is the most potent thrombin-specific inhibitor identified to date and is a representative pharmacologically active substance in leeches (Markwardt 1994; Chen et al. 2021). In the Pharmacopoeia of the People’s Republic of China (PPRC), antithrombin activity is the only standard for determining the quality of “Shuizhi” (Chinese Pharmacopoeia Commission 2020). It was not surprising that the extracts of *W. pigra* samples had antithrombin activities, as there were seven hirudins, of which at least one was shown to be functionally active (Liu et al. 2024). Although no hirudin was found in their genomes, *W. acranulata* (Li et al. 1997; Wang et al. 2013) and *W. laevis* (Li et al. 1997) were reported to have antithrombin activities, probably due to the antithrombin activity of other proteins such as the granulins (Hong and Kang 1999). The current version of the PPRC listed *W. pigra* and *W. acranulata*, but not *W. laevis*, as a legal drug material. However, since there were still dozens of antithrombic genes in *W. laevis*, we suggest that this species may also have application values for antithrombotic drug development in the future.

In conclusion, in this study we provide two nearly complete high quality genomes of *W. acranulata* and *W. laevis*. We identified many antithrombic gene families involved in anticoagulation, anti-platelet aggregation, fibrinolysis, and drug diffusion. We also estimated the relative expression levels of the antithrombic genes based on RNA-Seq data. Combined with our previous results from *W. pigra* and as well as *H. manillensis*, we systematically analyzed the similarities and differences on the genomes and especially their antithrombotic genes among the three non-hematophagous *Whitmania* leeches. This is the most comprehensive collection of genomes and leech antithrombotic biomacromolecules for the *Whitmania* leeches to date. Our results will greatly facilitate the evolutionary research and application of leech derivatives for medical and medicinal purposes of thrombosis.

## Materials and Methods

### DNA and RNA Sequencing

*W. laevis* and *W. acranulata* individuals were live-caught in Yutai, Shandong, China (GPS coordinates: E 116°28′55″, N 35°4′17″). Genomic DNA was extracted from fresh tissues using the DNeasy Blood and Tissue Kit (Qiagen, Germany) and was used to construct Nanopore and Illumina libraries. Total RNA was extracted from the head tissue using TRIzol reagent (Invitrogen), purified using the RNeasy Mini Kit (Qiagen, Chatsworth, CA, USA), and used to construct RNA-Seq (transcriptome) libraries.

For the Oxford Nanopore library preparation, a total of 10μg of high molecular weight DNA was subjected to random fragment shearing using a Megaruptor (Diagenode, NJ, USA). A long-read library (insert size, 20 kb) was constructed using the SQKLSK109 ligation kit and using the standard protocol. The purified library was loaded onto primed R9.4 Spot-On Flow Cells and sequenced using a PromethION sequencer (Oxford Nanopore Technologies, Oxford, UK) with 48-h runs at Wuhan Benagen Tech Solutions Company Limited, Wuhan, China. Base calling analysis of the raw data was performed using the Oxford Nanopore GUPPY software v0.3.0. The resulting ONT clean reads were used for genome assembly.

Similar to our previous study on *W. pigra* (Liu et al. 2024), one Hi-C, one Survey, and three RNA-Seq libraries for *W. laevis* and *W. acranulata* were constructed and were sequenced with 150 bp reads in both directions using the Illumina HiSeq 2000 sequencing platform. Quality control of raw reads for these three data was performed using fastp 0.20.0 with default settings and parameters (Chen et al. 2018).

### Genome assembling and repeat sequence finding

We used NextDenovo v2.5.0 (Hu et al. 2024) for de novo assembly of cleaned ONT reads and the ONT scaffolds were polished using NextPolish v1.4.0 (Hu et al. 2020). The polished sequences were further integrated with Hi-C reads using YaHS v1.1a (Zhou et al. 2023) and then imported into Juicebox v1.11.08 (Durand et al. 2016a) for Hi-C map visualization and manual optimization. Final pseudochromosome assemblies were generated using Juicer v1.6.2 (Durand et al. 2016b). We also used Survey reads to assemble the mitochondrial genome using GetOrganelle v1.7.7.0 (Jin et al. 2020).

To assess the thoroughness of the genome assembly, BUSCO v4.1.4 (Seppey et al. 2019) with the eukaryota_odb10 database was employed. Additionally, Merqury v1.3 (Rhie et al. 2020) was used to assess the quality of the assembly. RepeatModeler v2.0.3 and RepeatMasker v4.1.2-pl (Flynn et al. 2020) were used to search for repetitive sequences, and the repeat-masked scaffolds were used for gene prediction.

### Gene prediction and annotation

According to our recent study, a so-called BRAKER-plus strategy (Liu et al. 2023), which combines BRAKER prediction (Hoff et al. 2016) and our manual prediction, was used to identify antithrombotic genes. The GFF files from the process-oriented BRAKER prediction and the manual prediction were merged using AGAT v1.2.0 (Dainat 2021). After manua cleaning of duplicated features, we obtained a final version of the GFF file (BRAKER-plus.gff) with updated coordinate information for all potential antithrombotic genes. The coding sequences (CDS) and their corresponding protein sequences for all protein-coding genes were extracted using gffread v0.12.7 (Pertea and Pertea 2020).

### Phylogenomic and chromosome syntenic analysis

Combined with our previously reported genome of *W. pigra* (Liu et al. 2024), as well as genomic or transcriptomic data from eight other leech species, we tested the phylogenetic relationship among the three *Whitmania* species. The genome-derived CDSs of *H. medicinalis* (Kvist et al. 2020), *H. manillensis* (Liu et al. 2023), and *Dinobdella ferox* (Gao et al. 2023) were obtained directly from the gnome annotation results. The CDSs of the remaining five species were predicted from RNA-Seq data. The RNA-Seq reads of *Hirudo verbena* (SRR6926478), *Hirudo orientalis* (SRR6929415 ∼ SRR6929418), *Hirudinaria javanica* (SRR4162957), *Haemopis sanguisuga* (SRR10997447), *Haemadipsa cavatuses* (SRR4162952) were obtained from the Sequence Read Archive of NCBI, and were assembled using Trinity v2.9.0 (Grabherr et al. 2011). The CDSs were then predicted using GeneMarkS-T v5.1 (Tang et al. 2015). The orthologs of the CDS of all species were detected using OrthoFinder v2.3.11 (Emms and Kelly 2019), aligned using MACSE (Ranwez et al. 2011), and were concatenated using Seqkit v0.10.2 (Shen et al. 2016). IQ-TREE (Nguyen et al. 2015) was used to reconstruct trees using 1,000 bootstrap replicates with default settings.

We also used NGenomeSyn v1.41 (He et al. 2023) to visualize microcolinearity between chromosomes of the three *Whitmania* genomes. First, pairwise comparisons were performed using the GetTwoGenomeSyn.pl scripts with parameters set as “-MinLenA 5,000,000, - MinLenB 5,000,000, -MinAlnLen 5000, -MappingBin minimap2”. The results of the pairwise comparisons were then integrated using the main program NGenomeSyn.

### Expression analysis of antithrombotic genes

Using the RNA-Seq data sequenced in this study we estimated the expression characteristics of antithrombotic genes of *W. acranulata* and *W. laevis*. For each species, the CDS of all predicted genes, including the antithrombotic genes and the other non-antithrombotic genes, were used as mapping references. The salmon program (v1.0.0) (Patro et al. 2017) was used to calculate the transcripts per million (TPM) value for each gene based on the RNA-Seq data of each sample. To increase comparability, the TPM of genes from the same gene family were combined and the total TPM was calculated. The average TPM of all other non-antithrombotic genes was also calculated as background information. We also calculated the same values for *W. pigra* and *H. manillensis* using previously published RNA-Seq data (Liu et al. 2023; Liu et al. 2024). To ensure equal sample sizes, three RNA-Seq samples were randomly selected.

For each species, the TPM of each antithrombotic gene family was compared with the non-antithrombotic genes using the non-parametric two-related-samples test. The TPM of each gene family was compared between different samples using the non-parametric Mann-Whitney U test. In addition, to show the overall similarity of antithrombotic gene expression patterns between the two species, the gene expression spectra of all samples were clustered using hierarchical cluster analysis. All of the above statistical analyses were performed in SPSS v25.0 (IBM Corp., Armonk, NY, USA).

## Acknowledgements

The project was supported by the National Natural Science Foundation of China (No. 82260742 to GL), the Jiangxi “Double Thousand Plan” (No. jxsq2020101050 to GL and No. jxsq2023201063 to ZH), and the Science and Technology Foundation of Jiangxi Provincial Department of Education (No. GJJ190538 to ZH).

## Additional information

### Funding

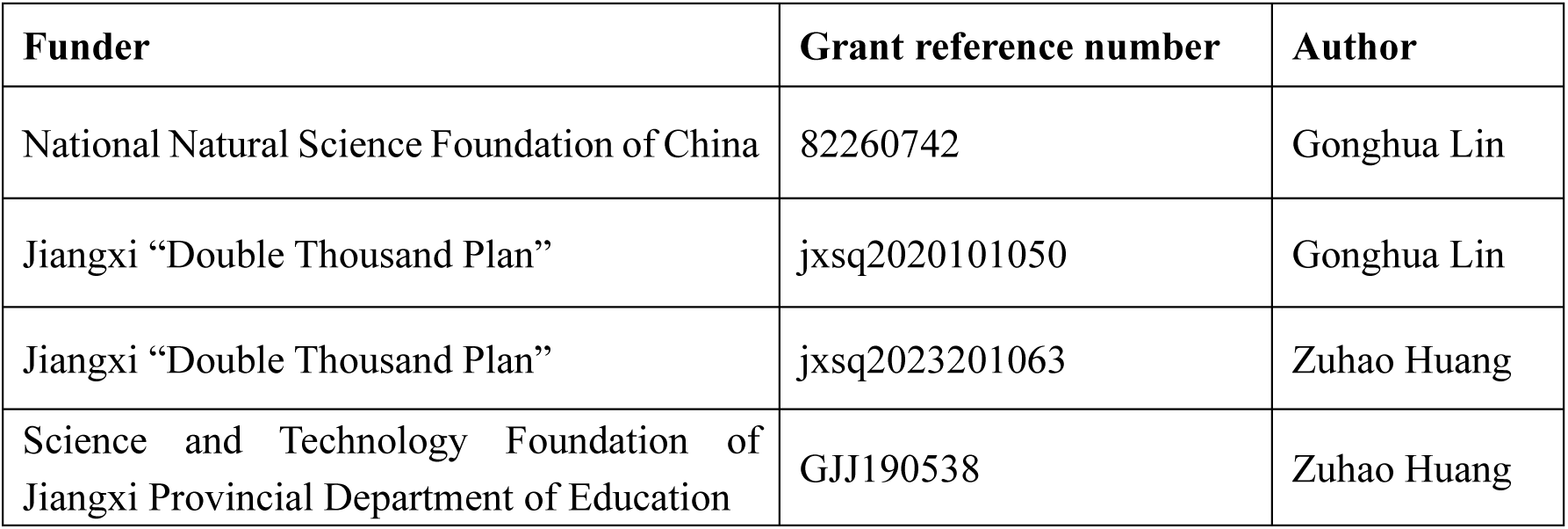

### Author contributions

Fang Zhao, Zichao Liu, and Gonghua Lin: Conceptualization, Data curation, Formal analysis, Writing – original draft & review & editing. Zuhao Huang: Investigation, Methodology, Writing – review & editing. Lizhou Tang and Bo He: Resources.

## Additional files

### Supplementary files

File_S1_Wacr.genome.zip: Supplementary File S1, the genome assembly of *W. acranulata*.

File_S2_Wlae.genome.zip: Supplementary File S2, the genome assembly of *W. laevis*.

File_S3_Wacr.gff.zip: Supplementary File S3, the GFF file of *W. acranulata*.

File_S4_Wlae.gff.zip: Supplementary File S4, the GFF file of *W. laevis*.

File_S5_Wacr.anti.fas: Supplementary File S5, coding sequences of all antithrombotic genes of *W. acranulata*.

File_S6_Wlae.anti.fas: Supplementary File S6, coding sequences of all antithrombotic genes of *W. laevis*.

These supplementary files were deposited in the Figshare online repository: https://doi.org/10.6084/m9.figshare.25757583

### Data availability

The raw data from our genome project was deposited in the SRA (Sequence Read Archive) database of National Center for Biotechnology Information with BioProject ID PRJNA1072149 and PRJNA1032729.

The following datasets were generated:

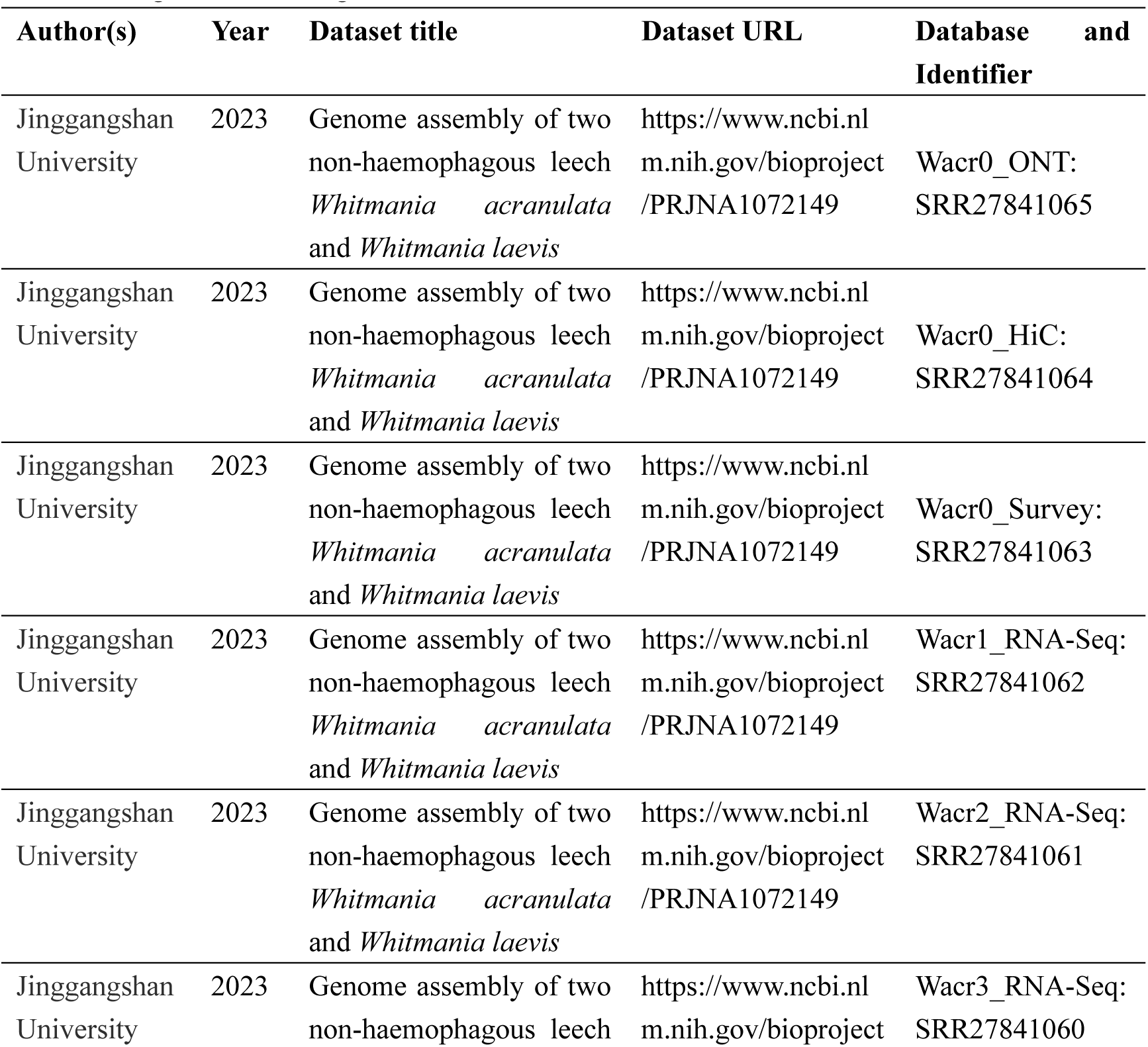
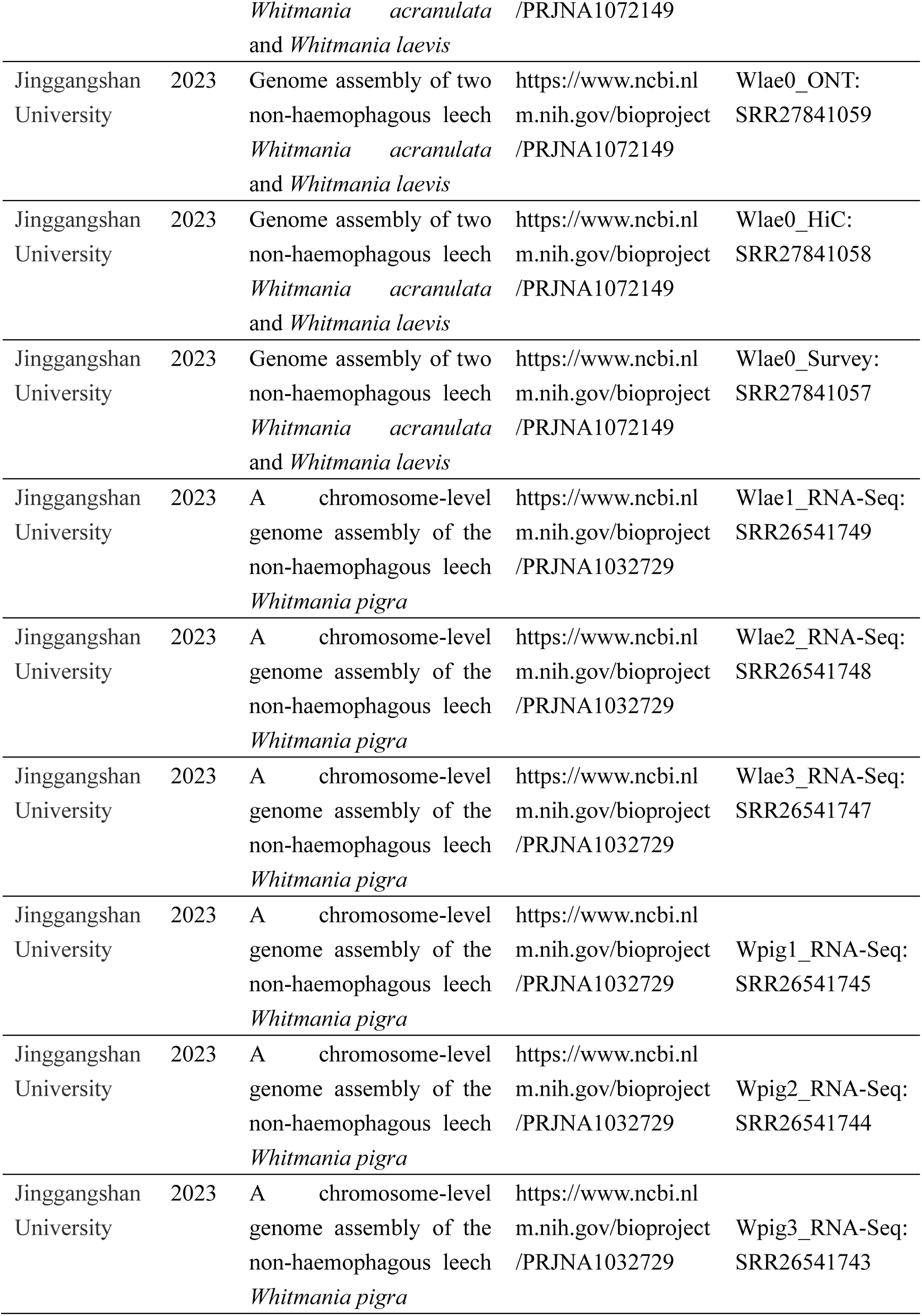

## Supplementary figures

Note: for each figure, the first protein sequence was the previously identified archetypal protein; the red triangles show conserved cysteines; the black stars mean stop codons; the red circle(s) indicate the reactive or binding residue(s) identified in the archetypal protein. For details, see our previous paper: Liu Z, Zhao F, Huang Z, He B, Liu K, Shi F, Zhao Z, Lin G. A chromosome-level genome assembly of the non-hematophagous leech *Whitmania pigra* (Whitman 1884): identification and expression analysis of antithrombotic genes. Genes, 2024, 15: 164.

**Figure S1.**
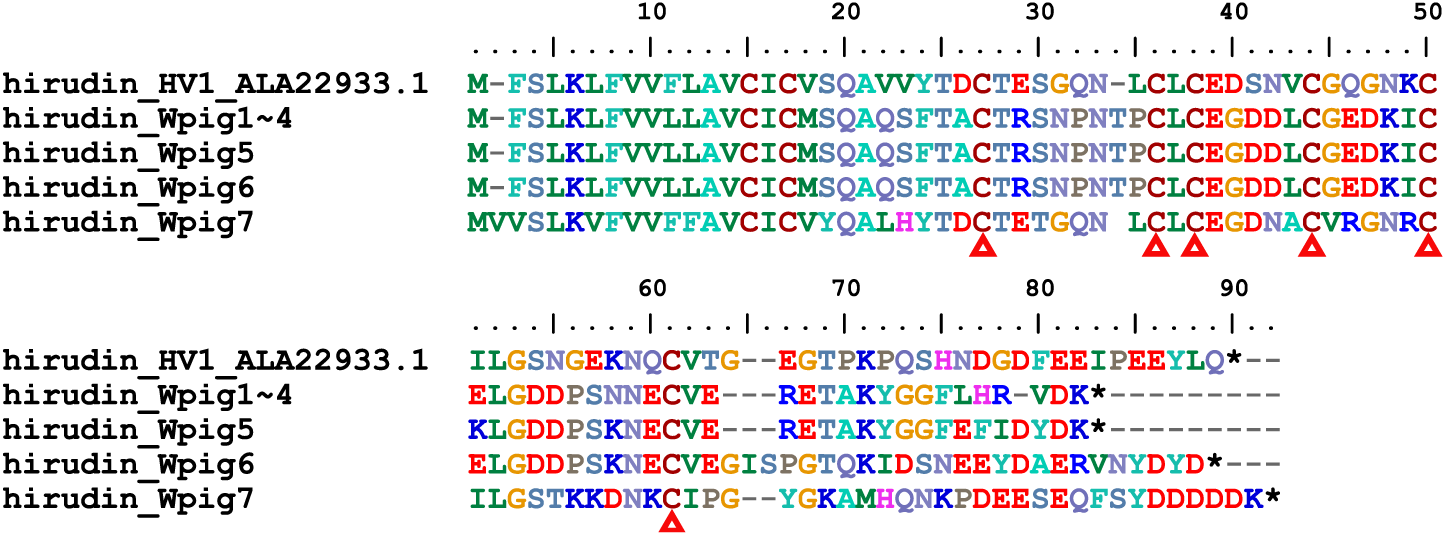
Alignment of archetypal hirudin and hirudins from *Whitmania pigra*.

**Figure S2.**
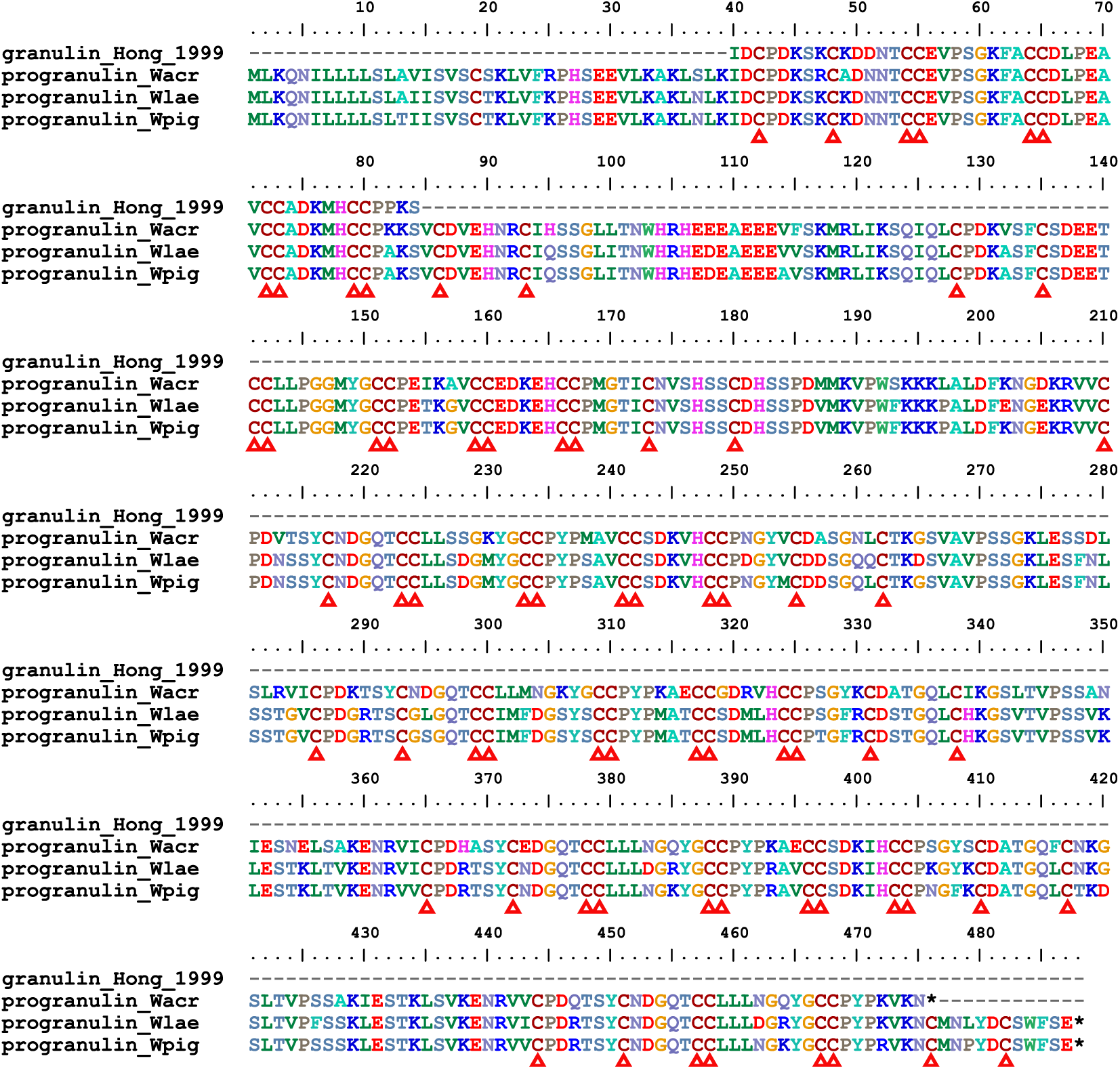
Alignment of archetypal granulin and progranulins identified from *Whitmania* species.

**Figure S3.**
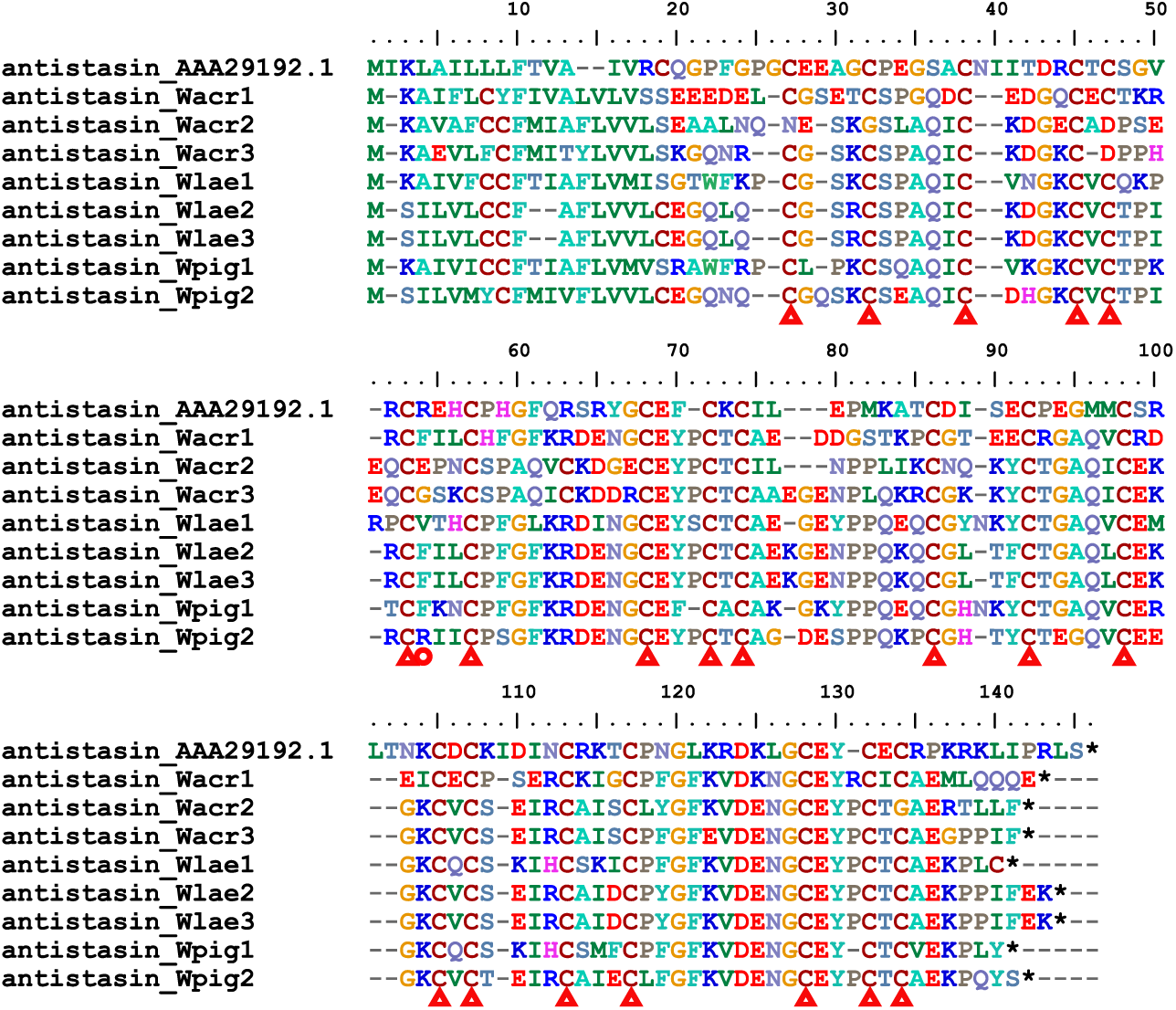
Alignment of antistasins.

**Figure S4.**
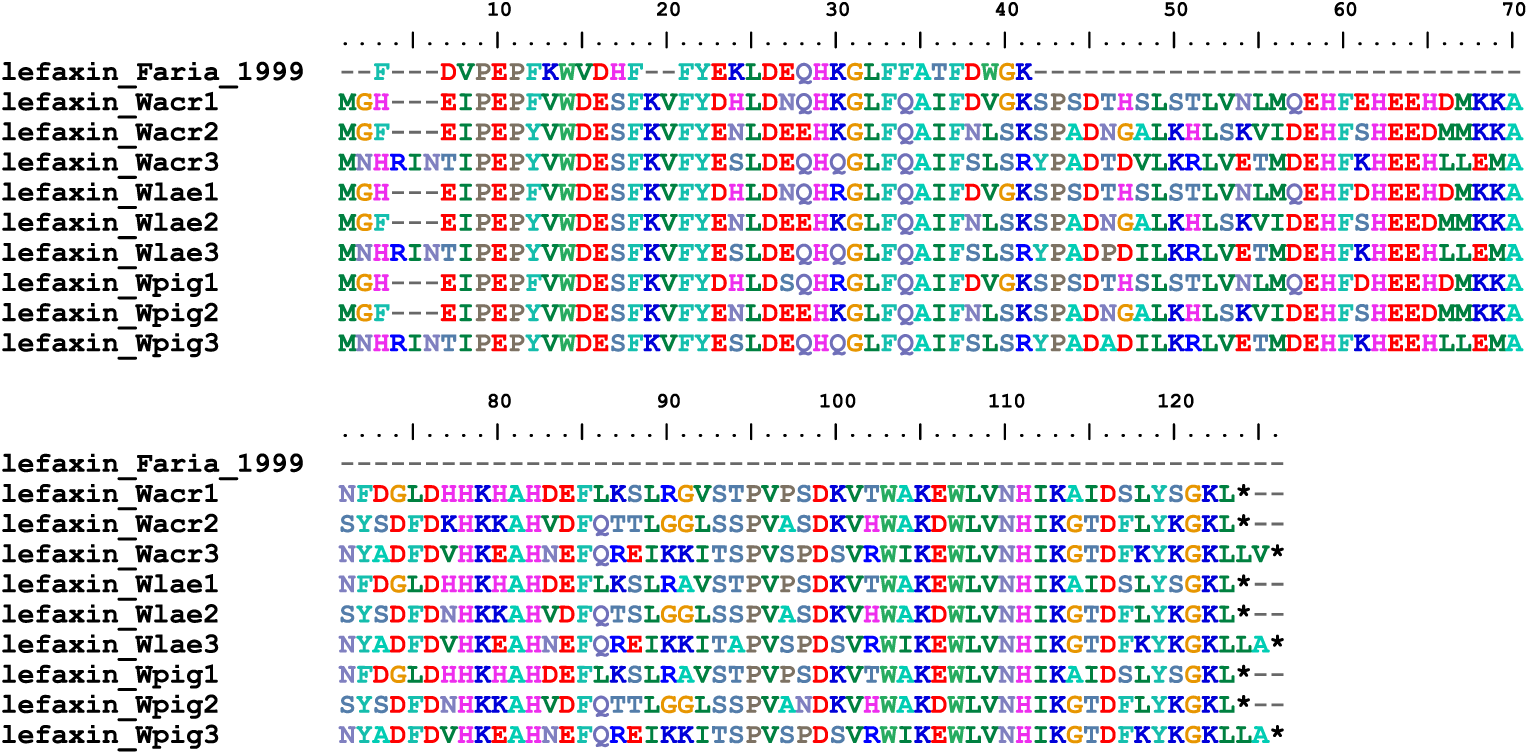
Alignment of lefaxins.

**Figure S5.**
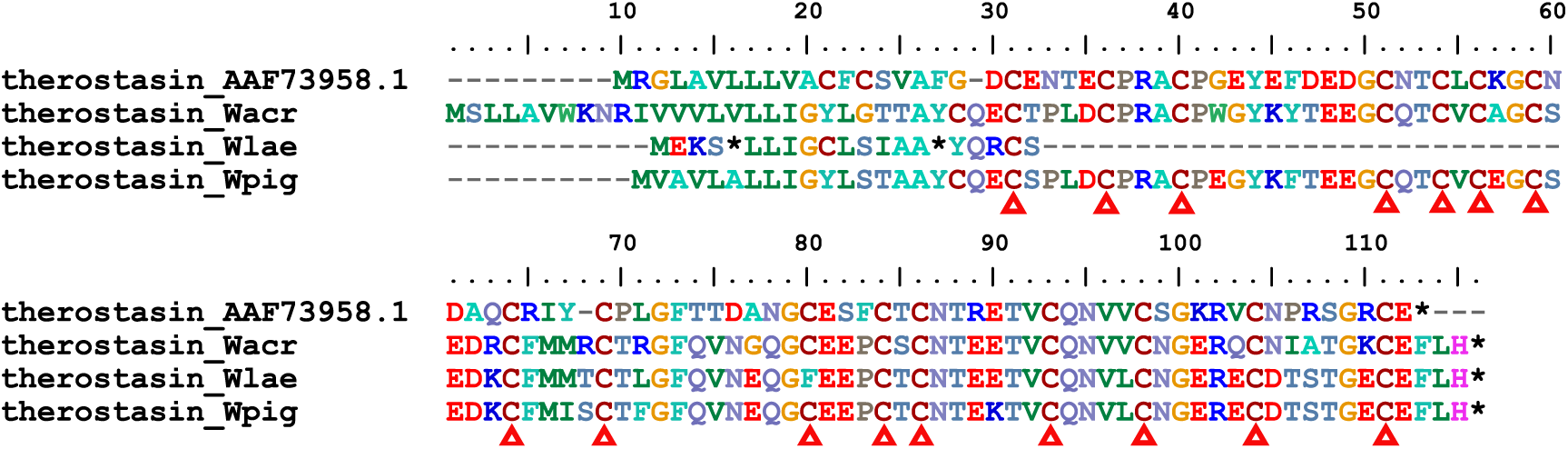
Alignment of therostasins.

**Figure S6.**
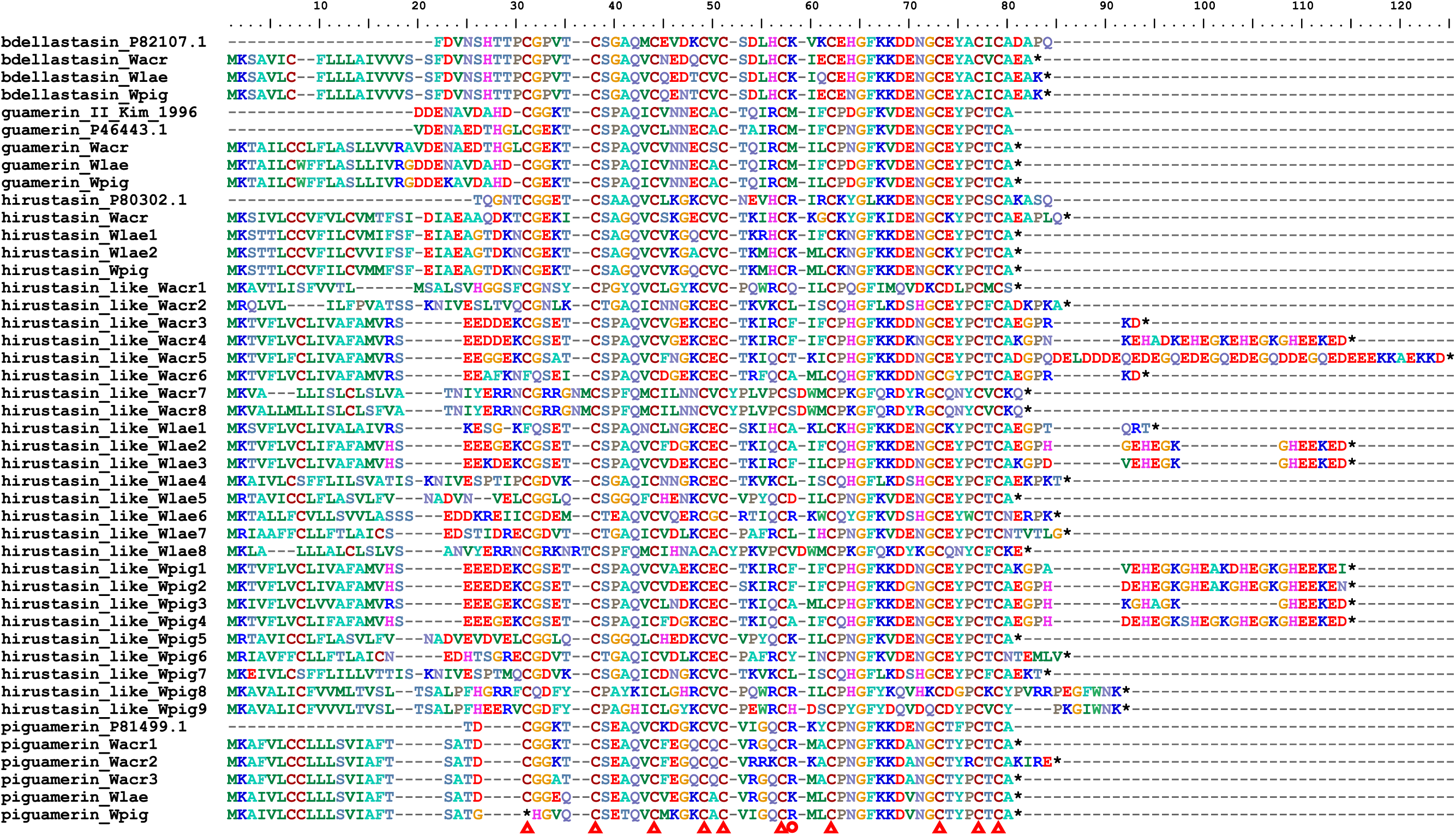
Alignment of proteins from hirustasin superfamily.

**Figure S7.**
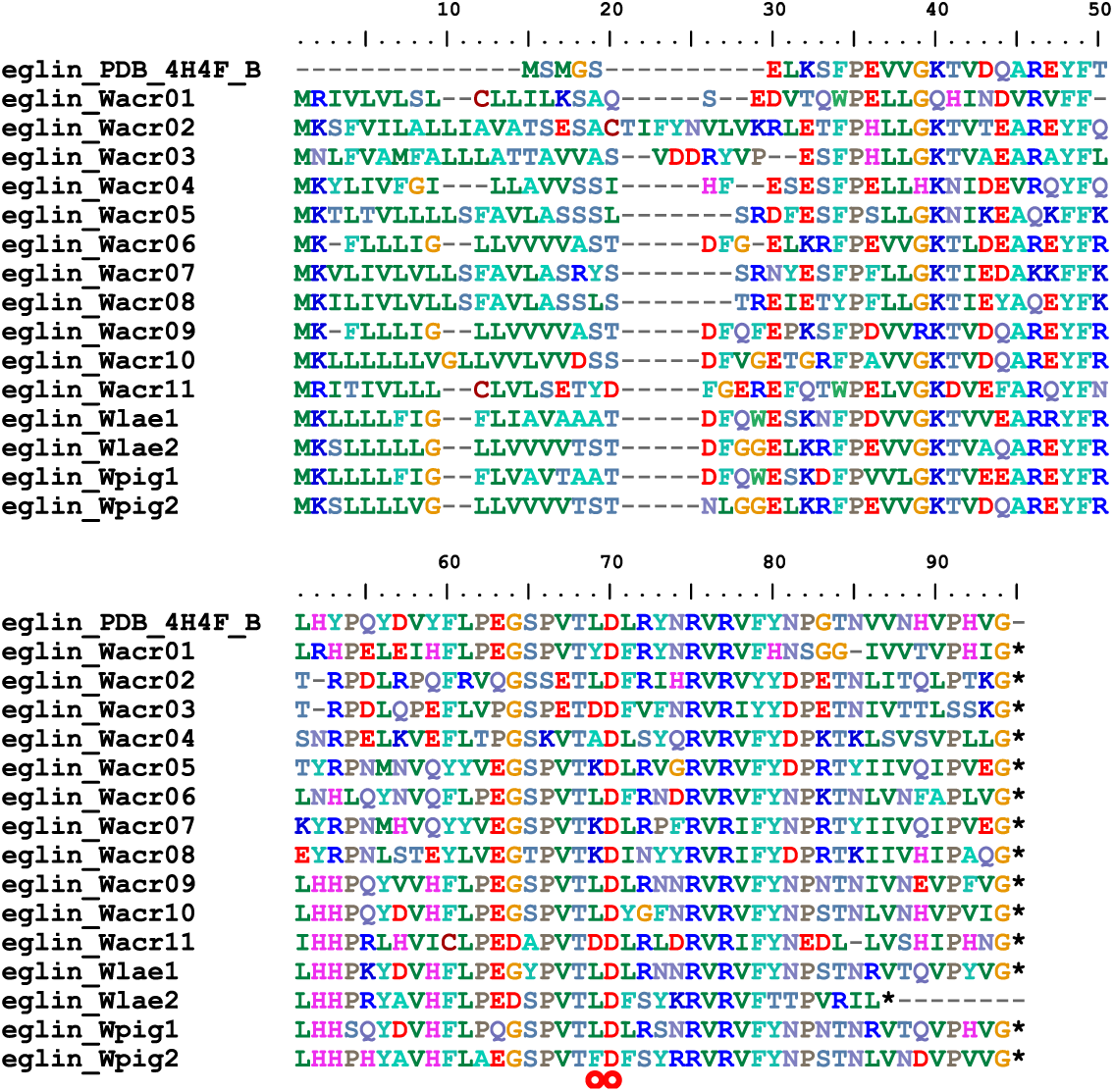
Alignment of eglins.

**Figure S8.**
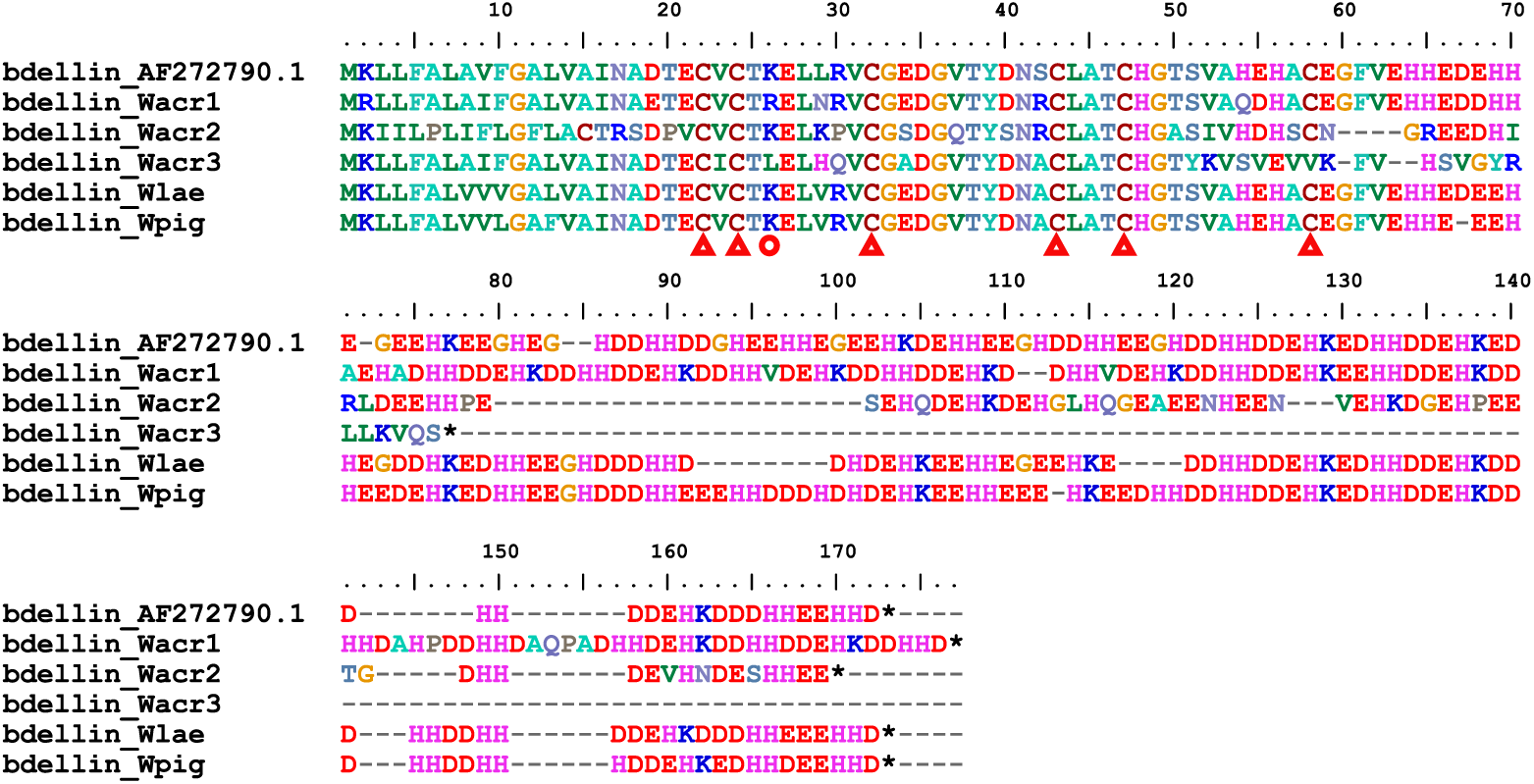
Alignment of bdellins.

**Figure S9.**
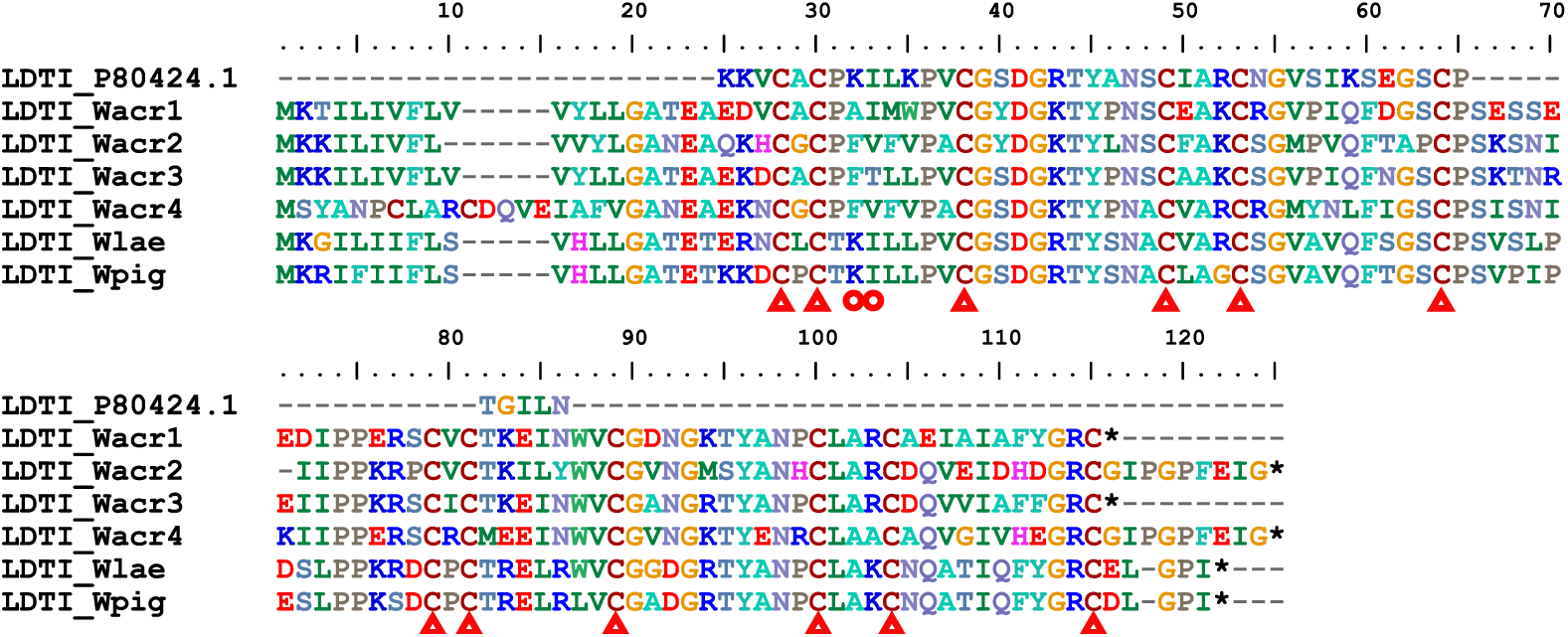
Alignment of LDTIs.

**Figure S10.**
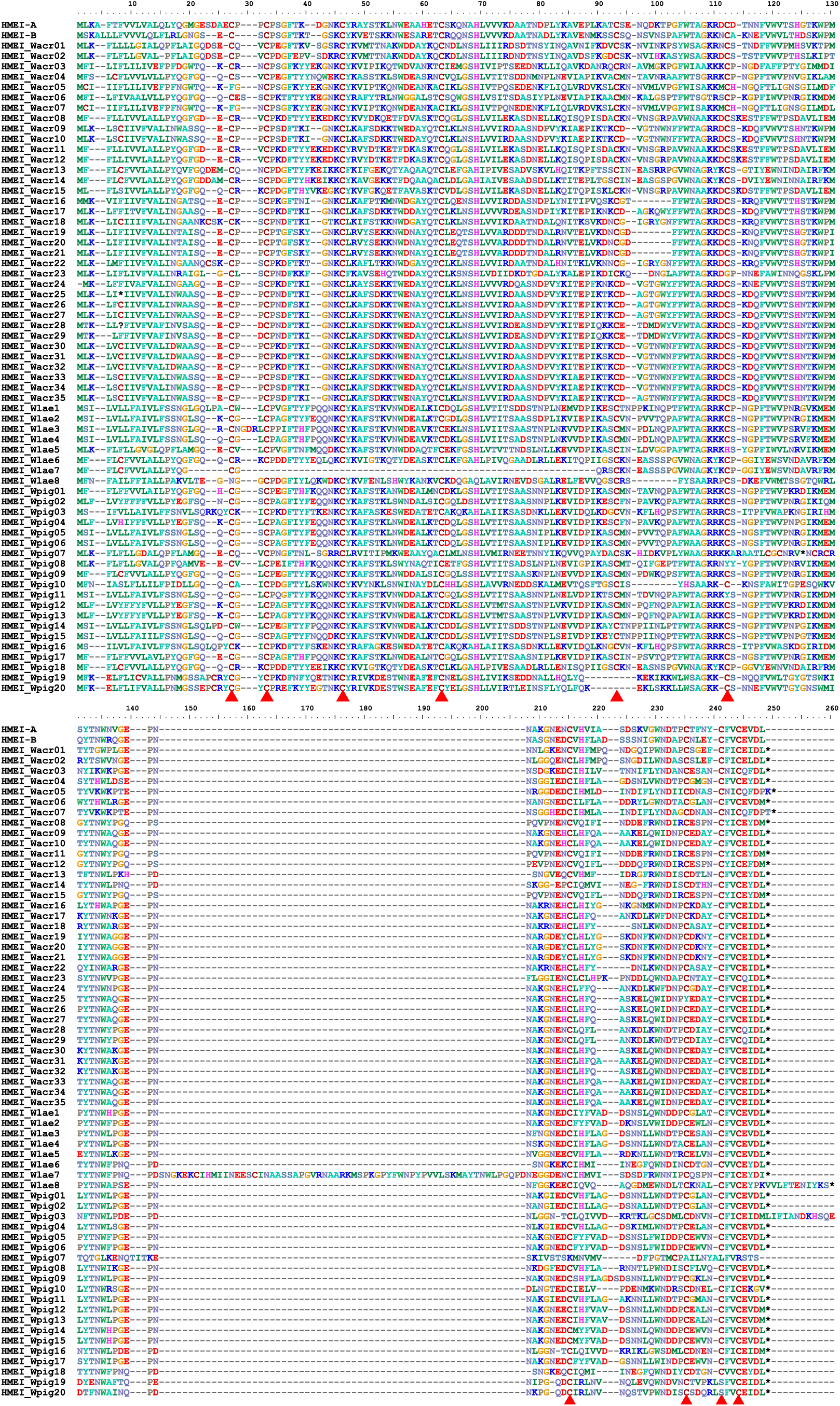
Alignment of HMEIs.

**Figure S11.**
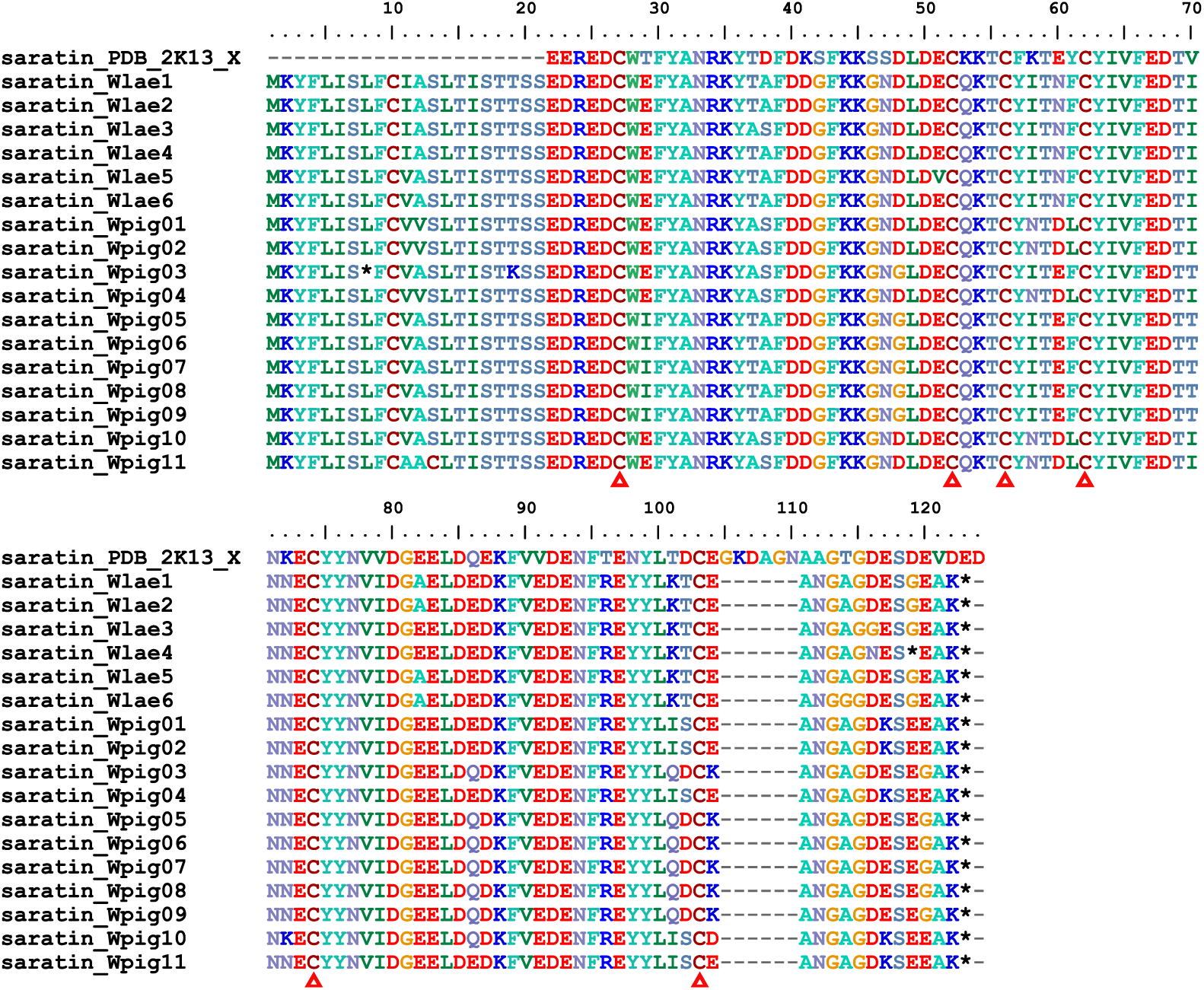
Alignment of saratins.

**Figure S12.**
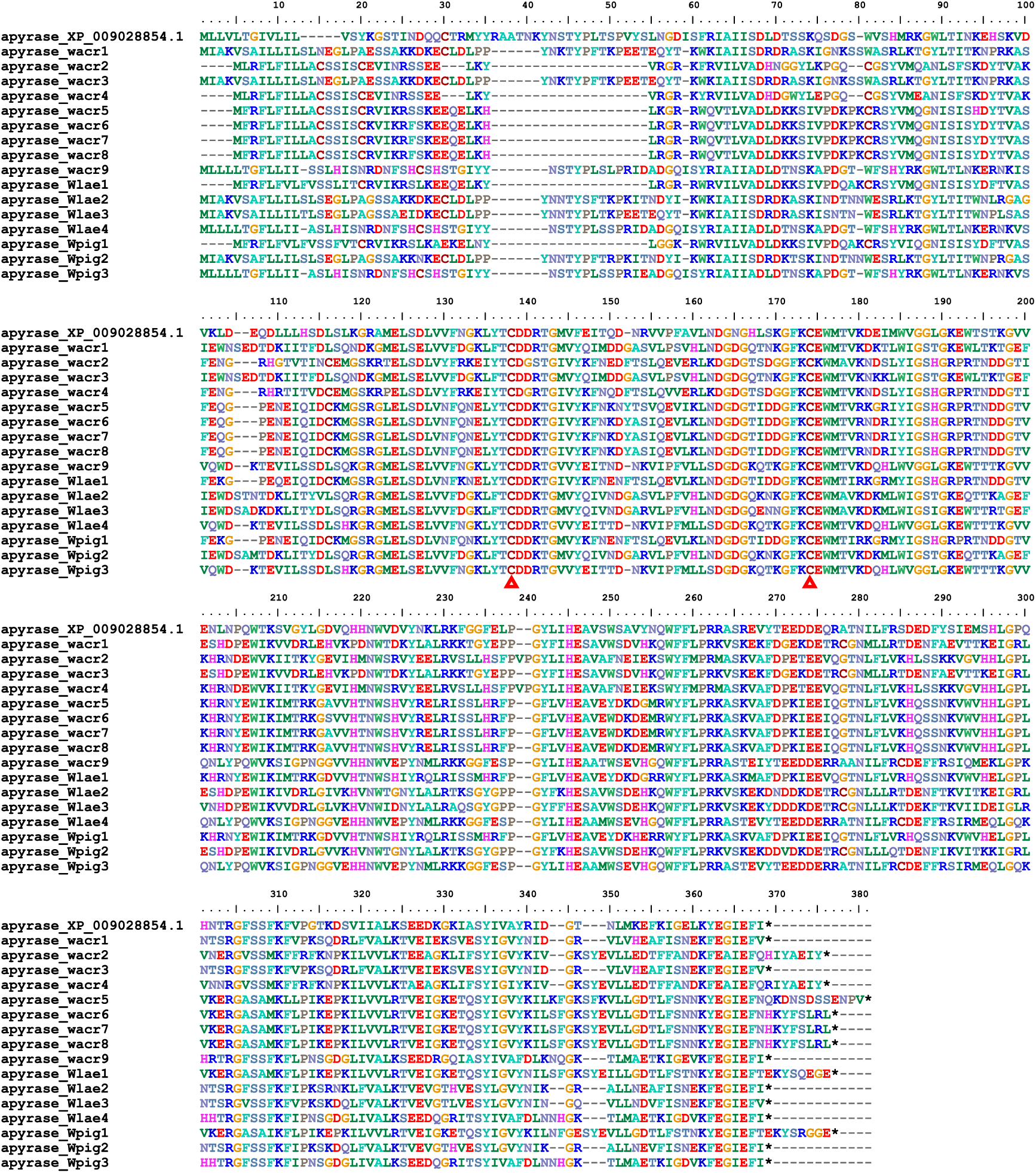
Alignment of apyrases.

**Figure S13.**
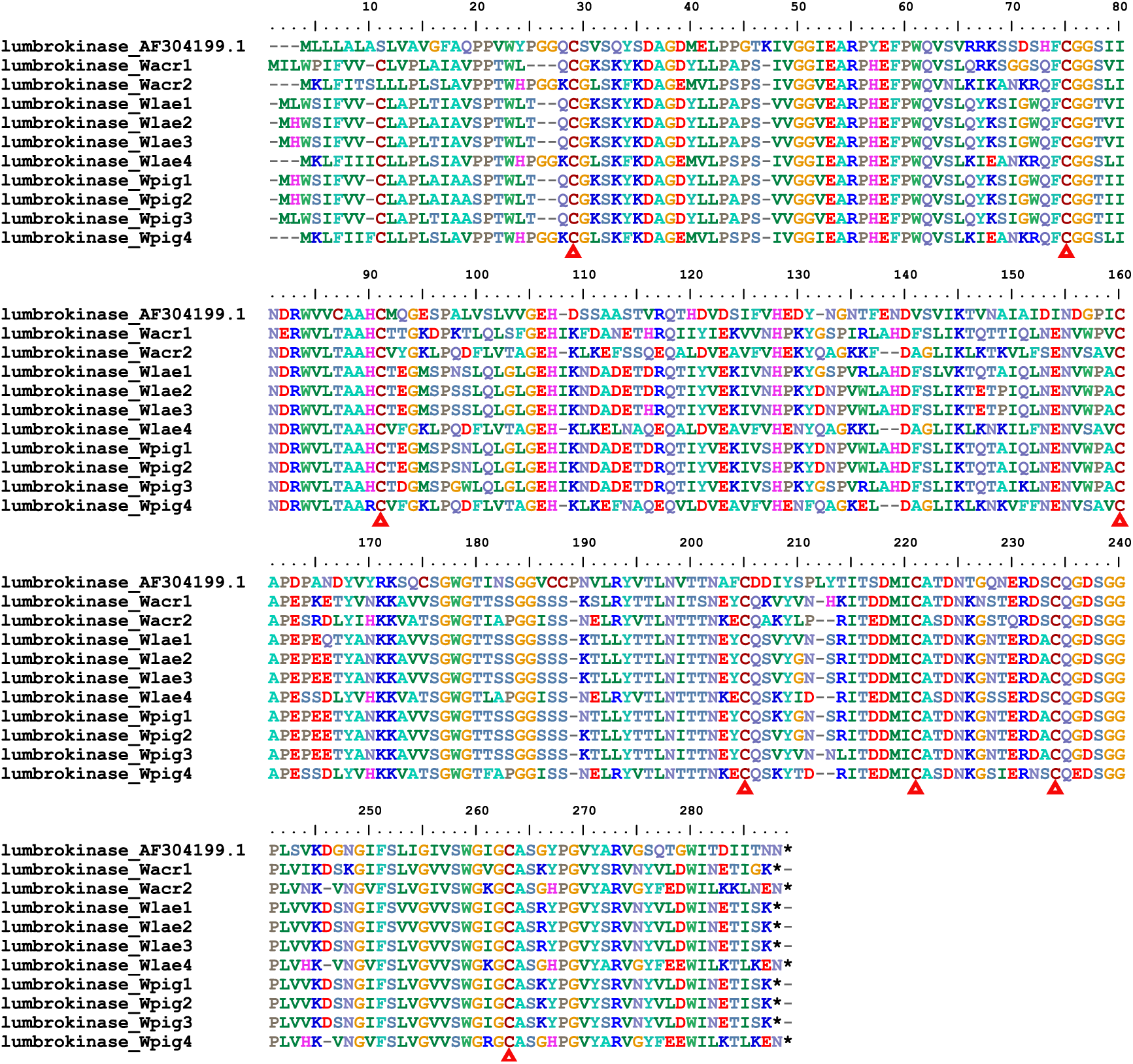
Alignment of lumbrokinases.

**Figure S14.**
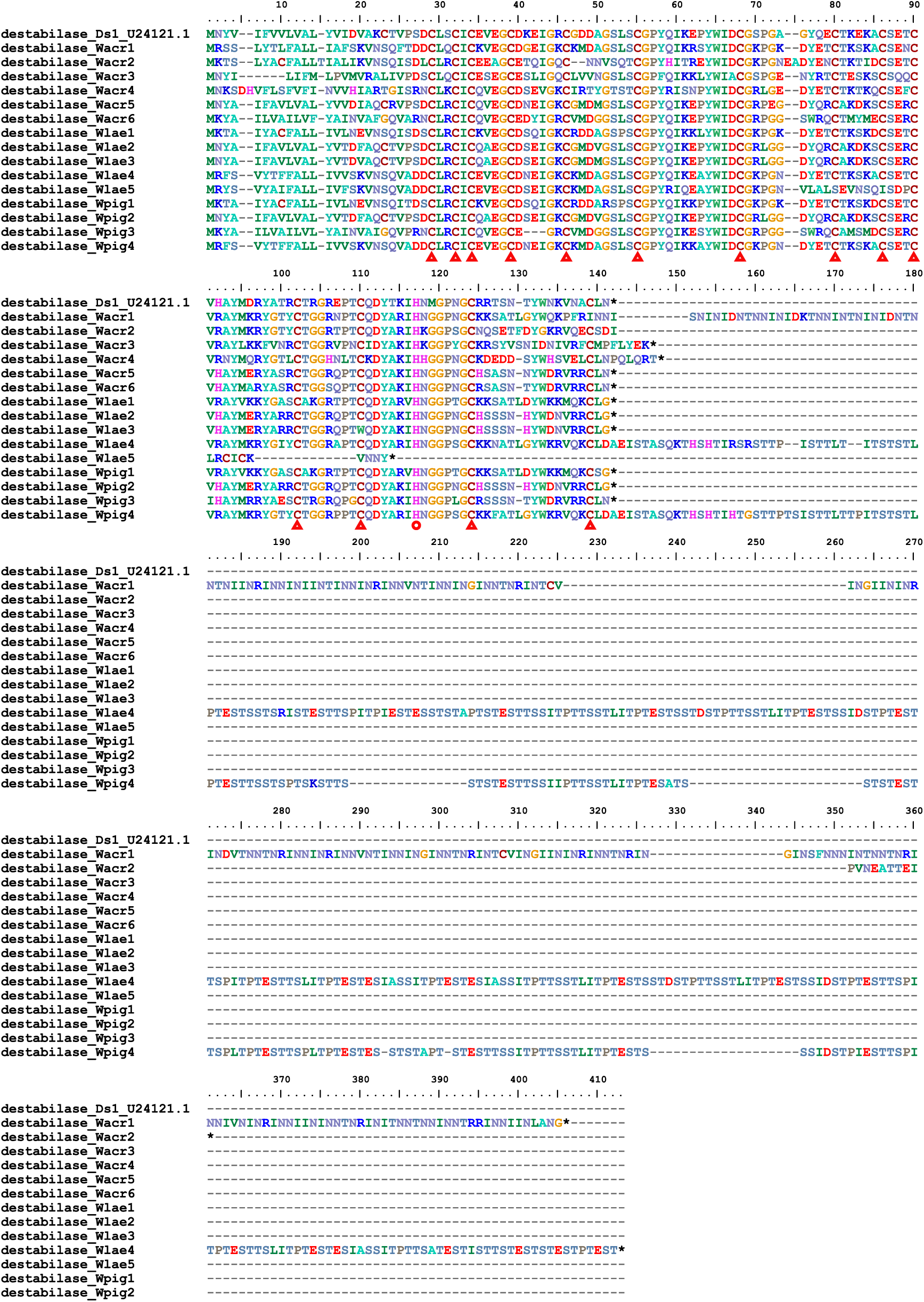
Alignment of destabilases.

**Figure S15.**
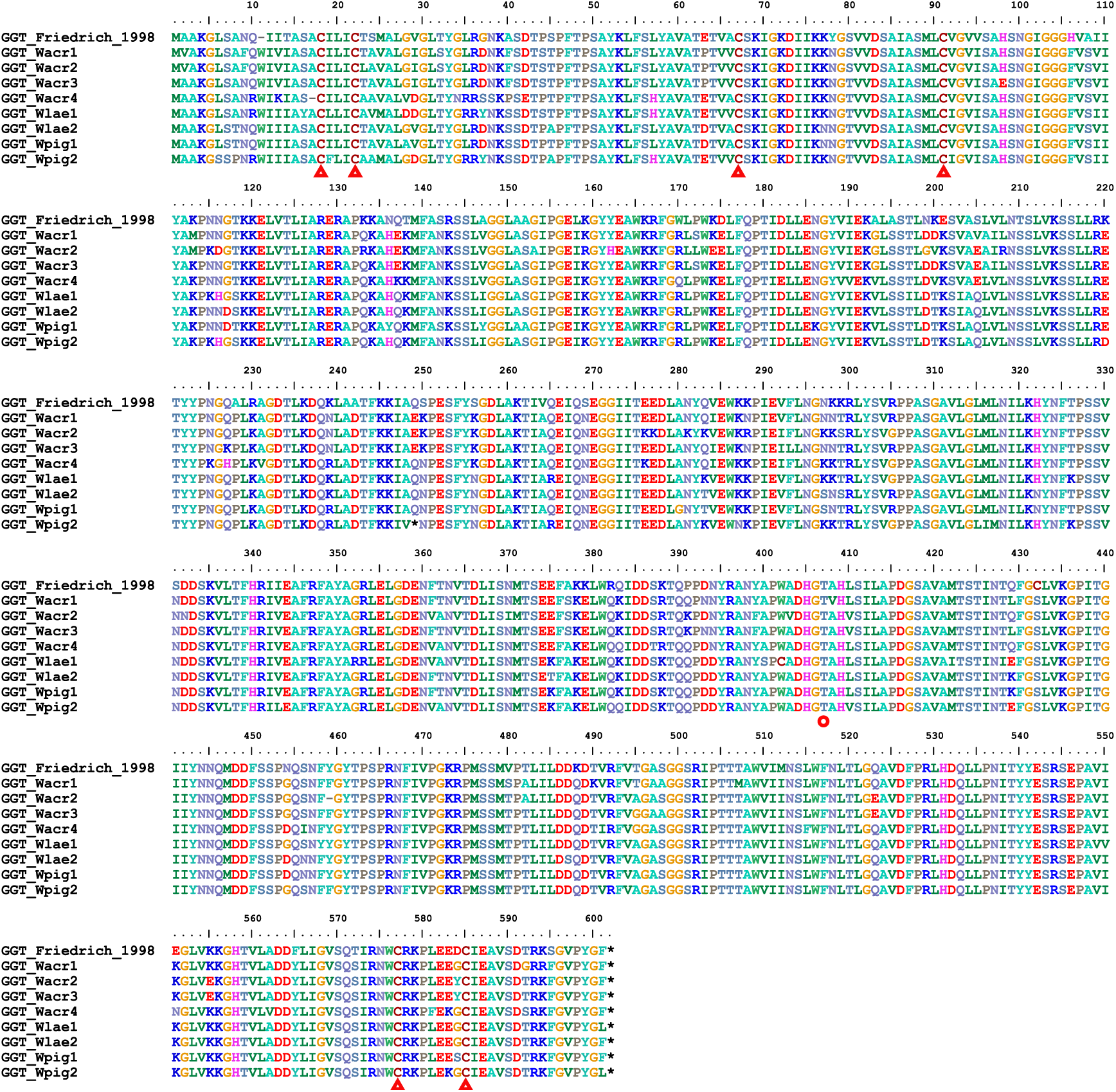
Alignment of GGTs.

**Figure S16.**
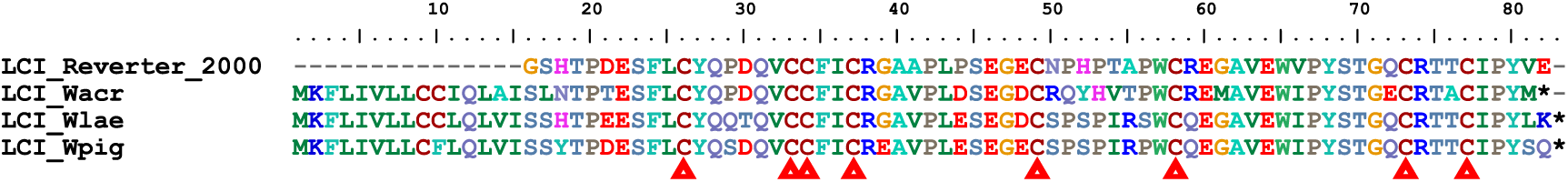
Alignment of LCIs.

**Figure S17.**
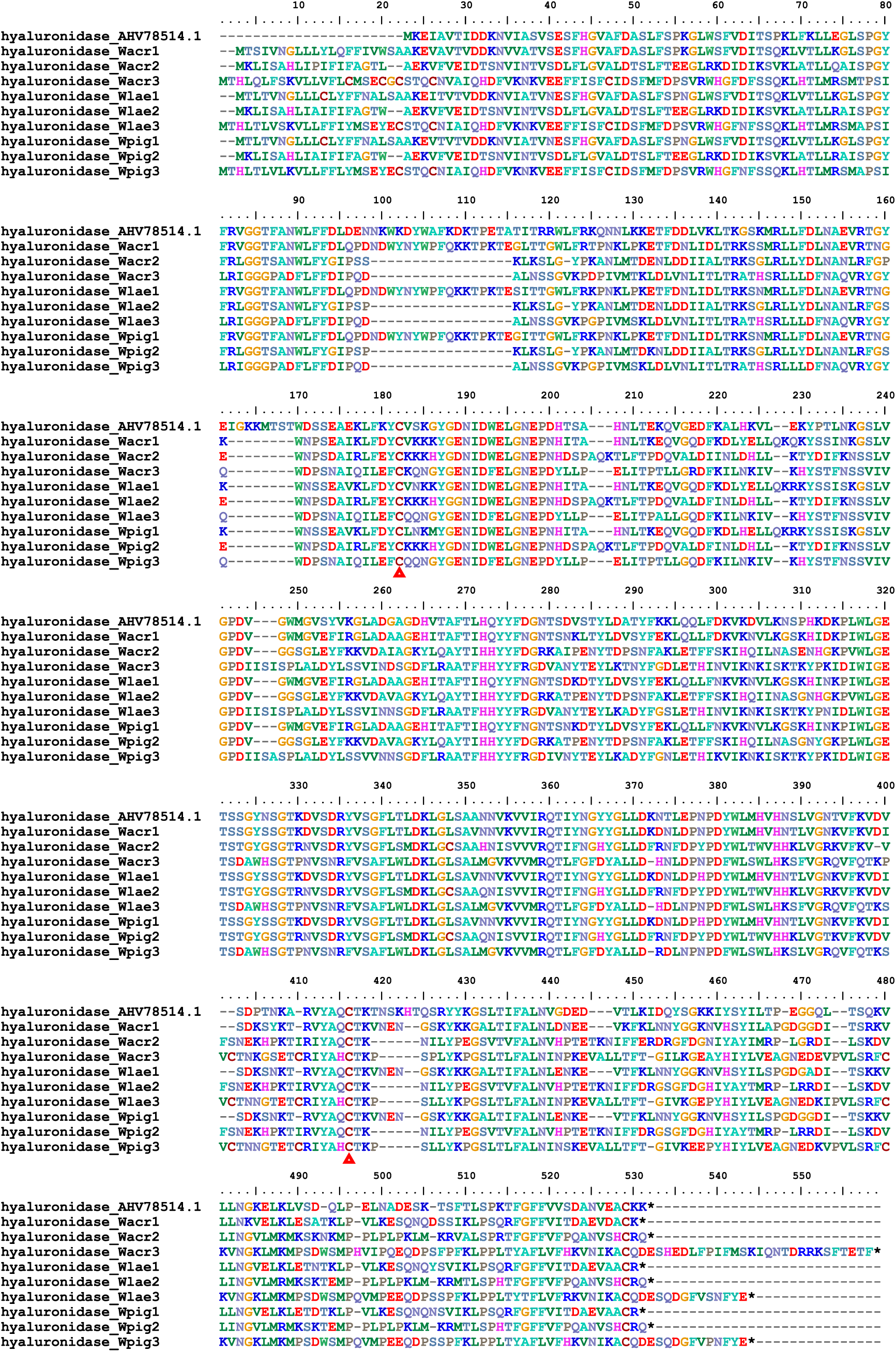
Alignment of hyaluronidases.

